# Spatial organization of voltage-gated ion channel expression across molecularly defined neuronal populations in the mouse mammillary bodies

**DOI:** 10.64898/2026.09.15.751911

**Authors:** Kristen S. Springer, Falyne C. Driver, Xiyue Wang, Heun Soh, Caleb Douala Moulema, William F. Flynn, Paul Robson, Anastasios V. Tzingounis, Alexander C. Jackson

## Abstract

The mammillary bodies (MB) are a hypothalamic component of the limbic Papez circuit that plays a critical role in spatial and episodic memory in mammals. Degeneration of the MB occurs in disorders associated with cognitive impairment, including Korsakoff’s syndrome and Alzheimer’s disease, yet the molecular organization and intrinsic properties of MB neurons remain poorly understood. Recent single-cell RNA sequencing identified multiple transcriptionally distinct neuronal populations within the MB and suggested that they differentially express voltage-gated ion channels that regulate neuronal excitability. Here, we used fluorescence *in situ* hybridization (FISH) to define the anatomical organization of cluster-enriched molecular markers and determine the spatial distribution of transcripts encoding voltage-gated sodium (Na_V_), potassium (K_V_), and hyperpolarization-activated cyclic nucleotide-gated (HCN) channels among defined subregions of the mouse MB. We found that marker transcripts occupy characteristic but partially overlapping spatial domains that broadly correspond to classical anatomical subdivisions. In addition, several ion channel transcripts, including *Scn1a, Scn2a, Kcnq2, Kcnq3,* and *Hcn1*, exhibited distinct patterns of enrichment across molecularly defined neuronal populations and MB subregions. Multiplex FISH further revealed unexpected co-expression of *Scn1a* and *Scn2a* within *Pvalb*-enriched neuronal populations, while whole-cell recordings demonstrated distinct intrinsic firing properties of neurons in the lateral and medial mammillary nuclei. Together, these findings establish a molecular framework linking neuronal population identity with voltage-gated ion channel expression in the MB and provide a foundation for future studies investigating how cell type-specific differences in intrinsic excitability contribute to memory function and neurological disease.

## INTRODUCTION

The mammillary bodies (MB) are a phylogenetically conserved subregion of the vertebrate hypothalamus, forming a critical node within the limbic Papez circuit by linking hippocampal subicular outputs with the anterior thalamus and midbrain (Aggleton and Brown, 1999; Aggleton et al., 2010; Vann, 2010; Jankowski et al., 2013; Dillingham et al., 2015; Vann and Nelson, 2015; Bubb et al., 2017). The MB are essential for spatial memory in animal models and for both spatial and episodic memory in humans, as lesions to the MB and its output pathways result in severe cognitive deficits (Vann, 2010; Bubb et al., 2017; Vann and Aggleton, 2004; Vann and Aggleton, 2003; Tsivilis et al., 2008; Vann, 2013). Consistent with this role, the MB and its associated fiber tracts undergo degeneration in disorders characterized by cognitive decline, including Korsakoff’s syndrome and Alzheimer’s disease (Kril and Harper, 2012; Dillingham et al., 2015; Arts et al., 2021; Saper and German, 1987; Grossi et al., 1989; Callen et al., 2001; Copenhaver et al., 2006; Baloyannis et al., 2016). In mouse models of AD, early β-amyloid deposition within the MB is accompanied by altered neuronal excitability (Canter et al., 2019; Huang et al., 2023). Together, these findings establish the MB as an important component of memory circuits and suggest that altered intrinsic excitability may contribute to disease-associated dysfunction. However, little is known about how molecularly defined MB neuronal populations are organized anatomically or how differences in ion channel expression contribute to their intrinsic properties.

Our single-cell RNA sequencing (scRNAseq) revealed that the mouse MB comprise multiple transcriptionally distinct neuronal populations, each characterized by the enrichment of unique combinations of genes (Mickelsen et al., 2020). These transcriptomic populations occupy characteristic, but partially overlapping, spatial domains that broadly correspond to established anatomical subdivisions of the MB. For example, neurons enriched for *Pvalb* and *Tac2* transcripts are preferentially distributed within the lateral mammillary nucleus (LM), whereas neurons enriched for *Nts*, *Calb1*, *Gpr83*, and *Pvalb* occupy distinct domains within the medial mammillary nucleus (MM). Importantly, these marker transcripts are enriched rather than exclusively expressed within individual populations and therefore do not define discrete anatomical boundaries. The identification of these molecularly distinct neuronal populations is broadly consistent with early electrophysiological studies demonstrating differences in firing properties between LM and MM neurons (Llinás and Alonso, 1992; Alonso and Llinás, 1992). Together, these observations suggest that molecularly distinct MB neuronal populations possess specialized intrinsic membrane properties that are likely shaped by differential ion channel expression. However, the molecular basis of these differences remains poorly understood.

Voltage-gated ion channels are key determinants of intrinsic neuronal excitability, shaping action potential generation, repetitive firing, synaptic integration, and neurotransmitter release (Bean, 2007; Vacher et al., 2008; Biel et al., 2009; Jan and Jan, 2012; Catterall, 2023). Among these, voltage-gated sodium (Na_V_), potassium (K_V_), and hyperpolarization-activated cyclic nucleotide-gated (HCN) channels have particularly important roles in regulating membrane excitability. Na_V_ channels mediate action potential initiation and propagation, Kv channels shape membrane repolarization and repetitive firing, whereas HCN channels contribute to resting membrane potential, dendritic integration and rhythmic firing (Bean, 2007; Vacher et al., 2008; Biel et al., 2009; Jan and Jan, 2012; Catterall, 2023).

Mutations in genes encoding Na_V_, K_V_ and HCN channels are among the most common causes of neurodevelopmental disorders (NDDs), including intellectual disability, autism spectrum disorder, Dravet syndrome and developmental and epileptic encephalopathies (Mantegazza et al., 2021; Singh et al., 1998; Abidi et al., 2015; Nappi et al., 2020; Marini et al., 2018; Kessi et al., 2022). Although genetically heterogeneous, many NDDs converge on altered neuronal excitability as a common pathophysiological mechanism and are frequently associated with impairments in learning and memory (Wang et al., 2020; Thapar et al., 2017; Ismail and Shapiro, 2019). These phenotypes are generally attributed to dysfunction of hippocampal and cortical circuits (Thapar et al, 2017; Ismail and Shapiro, 2019), whereas the potential contributions of other memory-related subcortical structures remain largely unexplored.

Despite increasing appreciation of the cellular diversity of the MB, little is known about how molecularly defined neuronal populations differ in the expression of ion channels that shape their intrinsic physiological properties. To address this question, we used scRNAseq-guided fluorescence *in situ* hybridization to map the spatial organization of molecularly defined MB neuronal populations and characterize the expression of voltage-gated ion channels that are critical in shaping intrinsic excitability. We found that cell type-specific markers define distinct MB subregions along the rostrocaudal axis and that several ion channel transcripts, including *Scn1a, Scn2a, Kcnq2, Kcnq3* and *Hcn1*, show subregion- and cell type-specific patterns of expression. Multiplex FISH revealed unexpected co-expression of *Scn1a* and *Scn2a* in *Pvalb*+ MB neurons, while slice electrophysiology demonstrated distinct intrinsic firing properties of LM and MnM neurons. Together, these findings define the molecular organization of MB neuronal populations, establish the spatial distribution of key voltage-gated ion channels implicated in NDDs, and provide a framework for understanding how population-specific ion channel expression may shape the intrinsic properties of MB neurons.

## MATERIALS AND METHODS

### Ethics statement

All experiments were performed according to the guidelines described in the National Institutes of Health Guide for the Care and Use of Laboratory Animals and were approved by the Institutional Animal Care and Use Committee of the University of Connecticut.

### Animals

For fluorescent *in situ* hybridization (FISH) experiments, P30-P35 male and female C57BL/6 (JAX stock #000664) mice were used. Immunohistochemistry and *in vitro* slice electrophysiology experiments were carried out using both C57BL/6 mice as well as parvalbumin (*Pvalb*)-Cre transgenic mice (*Pvalb*-IRES-Cre, JAX stock #017320), which were crossed to a Cre recombinase (Cre)-dependent tdTomato (tdT) reporter line (Ai14, JAX stock #007908) to visualize *Pvalb*+ neurons within subregions of the MB. The resulting cross is henceforth referred to as *Pvalb*-IRES-Cre;tdT. All mice were fed *ad libitum* and kept on a 12-hour light-dark cycle.

### Fluorescent *in situ* hybridization (FISH)

In preparation for FISH, male and female juvenile wild type C57BL/6 mice were anesthetized using isoflurane, decapitated, and brains were dissected out and frozen directly on dry ice. Brains were then embedded in cryo-embedding media (OCT) (Fisher Scientific, cat# 1437365) within cryomolds (Ted Pella, cat# 27112) on dry ice and stored at -80°C overnight or until experiments were performed. For experiments, 14 μm sections were taken and mounted directly onto SuperFrost Plus microscope slides (Fisher Scientific, cat# 50-949-342). After mounting, tissue sections were fixed using 4% paraformaldehyde (PFA), diluted from a 32% stock (Electron Microscopy Sciences cat#15714) at 4°C for 15 minutes, washed in 1x PBS, then dehydrated in increasing concentrations of ethanol (50, 70, and 100%) for five minutes each at room temperature. Dehydrated sections were stored in 100% ethanol at -20°C overnight before starting the FISH protocol.

The RNAscope Multiplex Fluorescent Assay V2 (Advanced Cell Diagnostics cat# 323110) was used according to the manufacturer’s protocol for single-plex FISH. Separate sets of serial sections were stained using probes targeting either *Calb1* (cat# 428431), *Gpr83* (cat# 317431), *Hcn1* (cat# 423651), *Hcn2* (cat# 427001), *Hcn3* (cat# 551521), *Hcn4* (cat# 421271), *Kcna2* (cat# 462811), *Kcnq2* (cat# 515141), *Kcnq3* (cat# 515151), *Nts* (cat# 420441), *Pvalb* (cat# 421931), *Scn1a* (cat# 556181), *Scn2a* (cat# 887241), *Scn8a* (cat# 434191), *Scn9a* (cat# 313341) or *Tac2* (cat# 446391). All probes used were designed and validated by Advanced Cell Diagnostics. The RNAscope Multiplex Fluorescent Assay V1 (Advanced Cell Diagnostics cat# 320851) was used according to manufacturer’s protocol for multiplex FISH. Sections were prepared for staining in the same way as the V2 kit, and the same probes were used. All sections collected for FISH were counterstained with DAPI and coverslips were placed after adding Prolong Gold Antifade Mountant (ThermoFisher Scientific cat# P36930).

### Image acquisition and analysis

Images (magnification of 40x and 100x) were acquired using a Leica SP8 confocal microscope. Images at 40x captured the entire MB, while 100x images of multiplex experiments were taken in central portions of each subregion. All images were processed using Fiji (Schindelin et al, 2012). All anatomical map overlays were acquired from the Paxinos and Franklin Mouse Brain Atlas (Paxinos, 2012) and adjusted to fit 40x ISH images based on a combination of the DAPI and mRNA signal.

Multiplex FISH images were analyzed using CellProfiler 3.1.9, collapsed 100x images were imported and a custom pipeline was used for analysis. A spreadsheet containing intensity values for each transcript generated by CellProfiler was then exported to an Excel spreadsheet for further analysis. A single composite image of each region with both cell type marker transcripts (either *Tac2*/*Pvalb* or *Nts*/*Calb1*) and the ion channel of interest (*Hcn1*, *Kcnq2*, *Kcnq3*, *Scn1a* or *Scn2a*) was used to set the region of interest (ROI) for intensity measurements. The maximum intensity value measured in a channel where no detectable transcript is expressed (intensity value of *Tac2* signal in the MnM for *Pvalb*/*Tac2* analysis and *Nts* signal in the LM for *Calb1*/*Nts* analysis) was used as the threshold, where intensity values above this number were considered positive expression and those below were considered negative.

### *In vitro* slice electrophysiology

Prior to recording, male and female juvenile C57BL/6 or *Pvalb*-IRES-Cre;tdT (P30-35) mice were deeply anesthetized with isoflurane before being transcardially perfused with ice-cold cutting solution containing the following (in mM): 7 NaCl, 75 sucrose, 25 glucose, 25 NaHCO3, 7.5 MgCl2, 2.5 KCl, 1.25 NaH2PO4, 0.5 CaCl2, and 5 ascorbic acid. Mice were then rapidly decapitated, their brains were removed and then cut into 225 μm slices using a vibrating microtome (V1 1200S, Leica). Slices were incubated at 36°C for 30 minutes in artificial cerebrospinal fluid (ACSF) containing the following (in mM): 125 NaCl, 25 NaHCO3, 11 glucose, 2.5 KCl, 1.25 NaH2PO4, 1 MgCl2, 2 CaCl2 and 0.1% biocytin. Tissue was then allowed to incubate at room temperature for an hour before recording. All solutions in which slices were incubated were continuously bubbled with 95%O2/5%CO2. Recordings were performed on both td+ and td-cells within the LM and MnM subregions of the mammillary bodies. Recorded cells were biocytin-filled and then subsequently stained.

### Biocytin staining and imaging

To stain filled cells, recorded slices were fixed in 4% PFA for 30 minutes, then moved to a solution of 0.5% Triton in PBS and stored at 4°C overnight. Slices were incubated in a solution of 0.5% Triton and 5% DNS in PBS with Alexa-488 conjugated streptavidin (1:500; ThermoFisher Scientific cat# S11223) for 2 hours then washed four times. Following the washes, slices were mounted onto slides with Prolong Gold Antifade Mountant with DAPI (ThermoFisher Scientific cat# P36931). Prepared slices were subsequently imaged on a Leica SP8 confocal.

### Slice electrophysiology data analysis

Electrophysiology data were analyzed using ClampFit (Molecular Devices) and Easy Electrophysiology software. Rheobase, Amplitude, 10-90% rise time, and 90-10% decay time were obtained using the action potential kinetics function in Easy Electrophysiology. Input resistance and fast afterhyperpolarization (fAHP) were calculated in ClampFit. Statistical comparisons between identified MnM and LM neurons were made using the Mann-Whitney test. For action potential count and input resistance, a two-way repeated-measure’s ANOVA was used to determine differences between neuronal subtypes at different current injections. Comparisons resulting in a p value less than 0.05 were considered statistically significant.

## RESULTS

### The MB are composed of several transcriptionally distinct subdivisions

Transcriptomic data from our scRNAseq analysis of the ventral posterior hypothalamus (VPH) from male and female C57BL/6 mice (Mickelsen et al., 2020) revealed six clusters with unique transcriptomic profiles that correspond to the MB (clusters 1-5 and 20) **(Fig. 1A and B)**. Using data from both the Allen Brain Atlas (Lein et al., 2007) and our own *in situ* hybridization for some of the most highly expressed genes in each cluster, we identified the approximate spatial domains occupied by these transcriptomic populations within the MB and their enrichment within specific subdomains of the MB **(Fig. 1C)**. These expression patterns are consistent with previous studies in both rat and mouse showing that parvalbumin (PV) immunoreactivity (IR) is concentrated in the dorsal median nucleus (dMnM), with additional expression in the LM. In contrast, calbindin (CB)-IR exhibits a largely complementary distribution, with staining throughout other subdivisions of the medial nucleus but little to no expression in the dMnM or LM. (Celio, 1990; Liu et al., 2025). From these data, it was evident that populations of neurons within each subdomain of the MB have unique gene expression profiles. We therefore sought to more precisely map the distribution of representative cluster-enriched transcripts across the rostrocaudal extent of the MB using fluorescence *in situ* hybridization (FISH).

**Figure 1:**
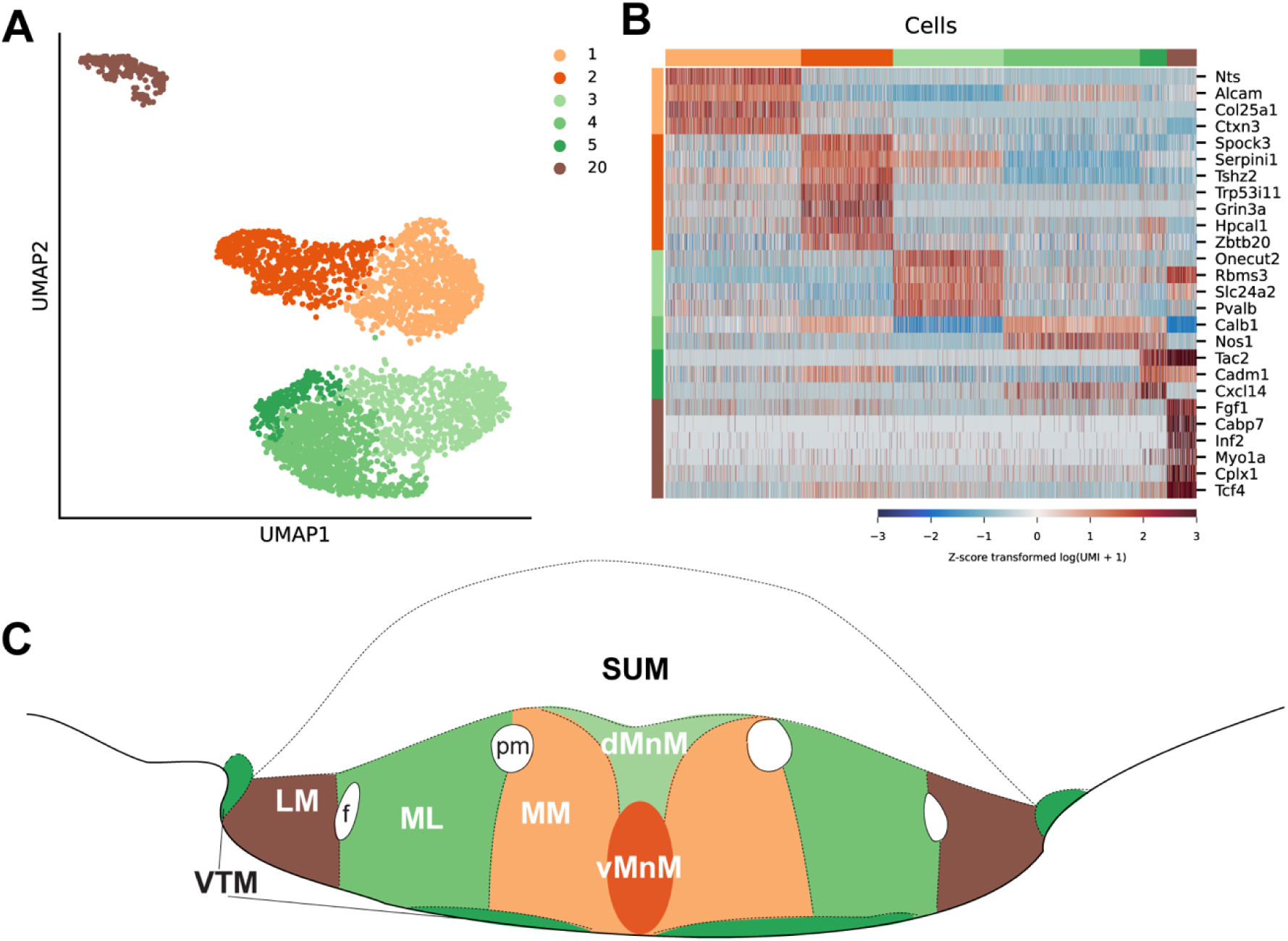
Transcriptomic markers map to distinct MB subregions. **A)** UMAP of the six clusters that comprise the MB (data from Mickelsen et al., 2020). **B)** Heatmap of the transcriptional markers of MB neurons within each subregion (using data from Mickelsen et al., 2020) **C)** Anatomical diagram of the major MB subdivisions based on the Paxinos and Franklin Mouse Brain Atlas (Paxinos, 2012), color-coded according to the transcriptomic clusters shown in A. Cluster-to-region assignments were based on the spatial expression patterns of cluster-enriched marker genes determined by FISH and the Allen Mouse Brain Atlas (ABA).

### Transcriptomic markers of MB subdivisions map to anatomically distinct subregions along the rostrocaudal axis

Although our previous scRNAseq analysis identified transcriptomic populations that map approximately to distinct spatial domains within the MB in a representative coronal section (Mickelsen et al., 2020), we sought to generate a higher-resolution anatomical map of the expression patterns of representative marker transcripts across the rostrocaudal extent of the MB (approximately Bregma −2.70 to −3.16 mm) **(Fig. 2A)**. To do this, we carried out single-plex FISH for these transcripts. More specifically, we probed for *Tac2*, *Pvalb*, *Calb1*, *Gpr83,* and *Nts* transcripts, each previously identified as enriched in subclusters representing the MB subdivisions. *Tac2* transcripts are abundant in the cluster representing subpopulations of LM neurons and FISH revealed that this is indeed the case. The anatomical data for *Tac2* also showed a pattern of enrichment in the ventral portion of the LM, with fewer cells being labeled in the dorsal portion. This observed pattern remained consistent throughout the MB **(Fig. 2Bi-v)**. Transcripts for *Pvalb* are mostly enriched in clusters predicted to represent LM and MnM neurons, and FISH revealed an enrichment of *Pvalb*-expressing neurons in the dorsal portion of the LM of the anterior-most sections **(Fig. 2Ci-ii)** that diminished more posteriorly **(Fig. 2Ciii-v)**. *Pvalb* signal in the MnM is detected in all sections, with most neurons being labeled in central MB sections. A small population of *Pvalb*+ neurons in the dorsal portion of the medio-lateral nucleus (ML) is also visible along the rostrocaudal axis.

**Figure 2:**
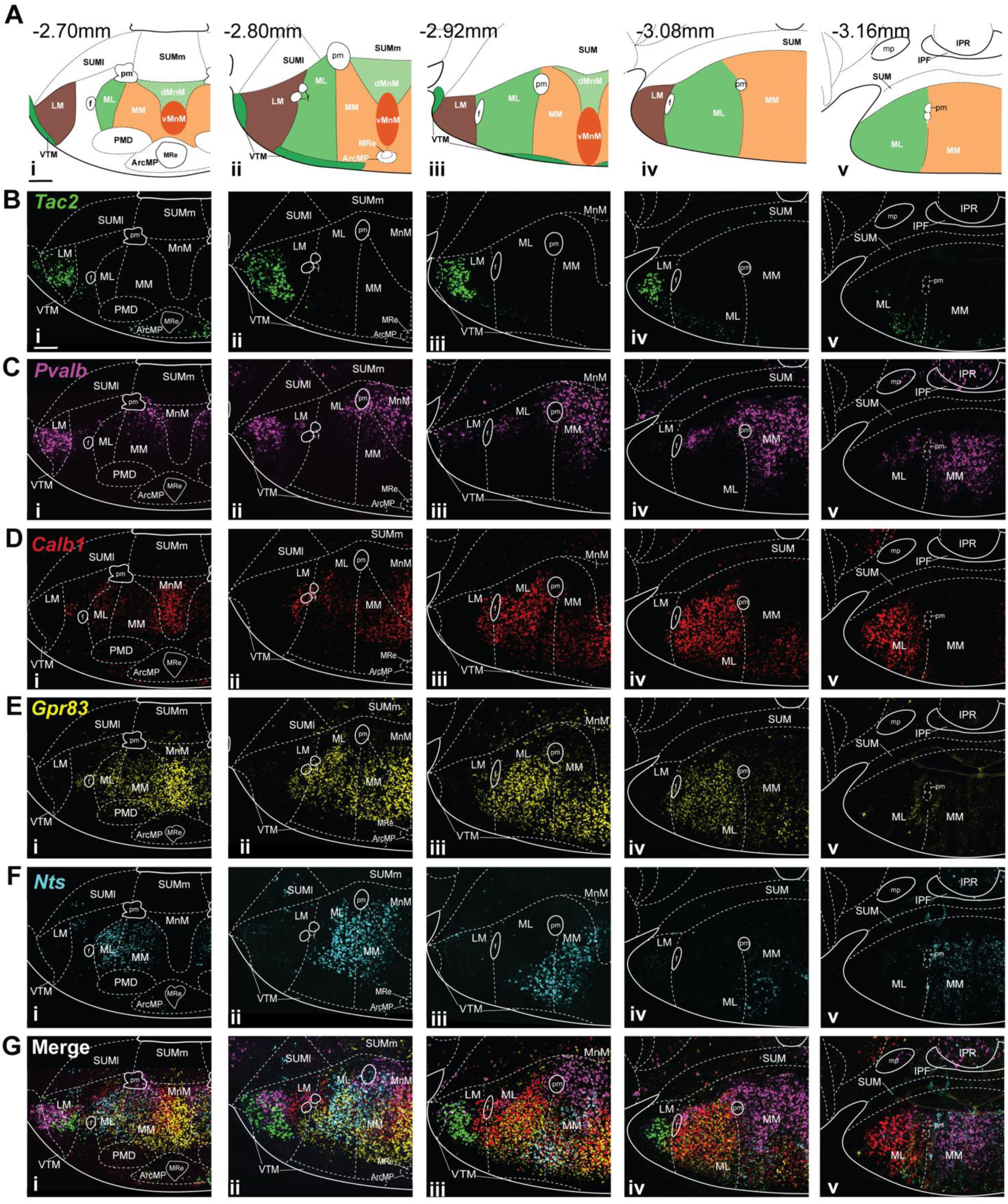
Anatomical map of transcriptomic markers of MB subregions. **A)** Representative maps of transcriptomic clusters to MB subregions based on the most highly expressed transcripts within these clusters. Map outlines are from the Paxinos and Franklin Mouse Brain Atlas (Paxinos, 2012) with their approximate Bregma coordinates indicated in the top left of each image going from anterior (i) to posterior (v) (scale bar=200 μm). These outlines were overlaid on the FISH images included in B-G. **B)** Confocal images (40x) of coronal sections taken from wildtype mice showing that *Tac2* expression is concentrated in the LM (n=4) (scale bar=200 μm). **C)** Representative images of FISH for *Pvalb* transcripts throughout the MB, *Pvalb*+ neurons are found in both the LM and multiple regions of the medial nucleus (n=3). **D)** Images showing the expression pattern of *Calb1* within MB subnuclei, these neurons are primarily in the ML (n=4). **E)** Images showing transcripts for *Gpr83* are strongly expressed in the medial nucleus of the MB diminishing in posterior sections (n=4) (iv-v). **F)** Confocal images of FISH for *Nts*-expressing neurons showing they are mostly found in the medial nucleus (n=3). **G)** Merged images of transcripts for each cell cluster-specific marker from anterior (i) to posterior (v) showing enrichment of each of these markers within the MB.

Our scRNAseq data identified *Calb1* as most highly expressed within the medial nucleus, with the greatest enrichment in the ML. Consistent with these findings, FISH revealed that *Calb1*-expressing neurons are concentrated within the MnM and ML in anterior sections, with expression becoming more prominent in ventral portions of the MM and MnM in progressively more posterior sections **(Fig. 2Di–v)**. *Gpr83* transcripts were also detected throughout several subdomains of the medial nucleus, with the strongest expression beginning in anterior sections and gradually diminishing along the rostrocaudal axis **(Fig. 2Ei–v)**. Finally, *Nts* transcripts were enriched within the MM-associated spatial domain, consistent with the scRNAseq data, and this pattern was maintained throughout the rostrocaudal extent of the MB **(Fig. 2Fi–v)**. However, we also observed a distinct population of *Nts*+ neurons extending into the ML, illustrating that individual marker transcripts define domains of enrichment rather than discrete atlas-defined anatomical subdivisions. Finally, composite images of the single-plex FISH data reveal the relative spatial organization and overlap of these marker-enriched domains **(Fig. 2G)**.

An important consideration in interpreting these findings in the context of MB anatomy is that transcriptomic clusters and atlas-defined anatomical subdivisions represent different levels of organization. In our previous scRNAseq study, neuronal clusters were identified from transcriptome-wide expression profiles, with marker genes subsequently selected based on their relative enrichment within individual clusters, where combinations of these markers could reliably predict cluster identity (Mickelsen et al., 2020). Mapping individual marker transcripts by FISH therefore identifies spatial domains in which neurons associated with a given transcriptomic population are enriched but does not establish a one-to-one correspondence among marker genes, transcriptomic populations, and anatomically defined MB subdivisions. We found that marker-enriched populations frequently overlap or extend across atlas-defined boundaries, and individual transcripts are rarely restricted to a single anatomical region. We therefore used boundaries from the Paxinos and Franklin mouse brain atlas (Paxinos, 2012) as an anatomical reference rather than as molecularly defined borders between MB subdivisions. Accordingly, throughout these analyses, descriptions of a transcript as enriched within a particular MB subregion indicate preferential expression within that approximate spatial domain rather than exclusive or uniform expression within the atlas-defined subdivision. This framework also guided our interpretation of ion channel transcript distributions relative to marker-enriched neuronal populations. Overall, these data provide a detailed anatomical map of cluster-enriched transcript expression and suggest that the molecular organization of the MB may refine, rather than precisely recapitulate, its classically defined anatomical subdivisions.

### Ion channels are differentially expressed throughout the VPH and MB

As a first step to characterizing MB neurons, we leveraged our scRNAseq data to develop a comprehensive compendium of ion channel transcript expression both throughout the entire VPH **(Fig. 3A)** and within the 6 clusters representing MB neurons specifically, clusters 1-5 and 20 **(Fig. 3B)**. Notably, several families of ion channels show a pattern of enrichment in the clusters representing MB subdivisions relative to those representing other portions of the VPH. These include members of the voltage-gated potassium (K_V_), sodium (Na_V_), and calcium (Ca_V_) channel families as well as the cyclic nucleotide-gated (CNG) channel family, all of which are key to determining the intrinsic firing properties of the neurons in which they are expressed. Transcripts for several K_V_ channels such as K_V_1.2 (*Kcna2*), K_V_3.1 (*Kcnc1*) and K_V_7.2 (*Kcnq2*) were detected across the VPH (Fig. 3A). Each channel plays a role in both shaping the action potential and preventing hyperexcitability (Jan and Jan, 2012). Among these channels, some patterns of enrichment begin to emerge. For example, while *Kcnq2* is evenly expressed across the VPH, K_V_7.3 (*Kcnq3*) transcripts are more strongly expressed in VPH cluster 20, representative of LM neurons. While also found elsewhere in the VPH, *Kcna2* is most strongly expressed in the 6 MB clusters, specifically in clusters 3 (dorsal MnM neurons) and 20. Our data further show that transcripts for the inwardly rectifying potassium channels *Kcnj3* and *Kcnj9* are enriched in the medial MB clusters relative to the rest of the VPH (Fig. 3A), with particularly strong expression in the dorsal MnM (Fig. 3B). Additionally, transcripts for the two-pore potassium channel *Kcnk9* are enriched within the MB, with strong expression in the LM subcluster (Fig. 3B). All three channels are associated with action potential duration and stabilizing resting membrane potential (Hallmann et al., 2000; Brickley et al., 2007; Hibino et al., 2010; Enyedi and Czirják, 2010) and may therefore contribute to functional differences between MB neuronal subpopulations.

**Figure 3:**
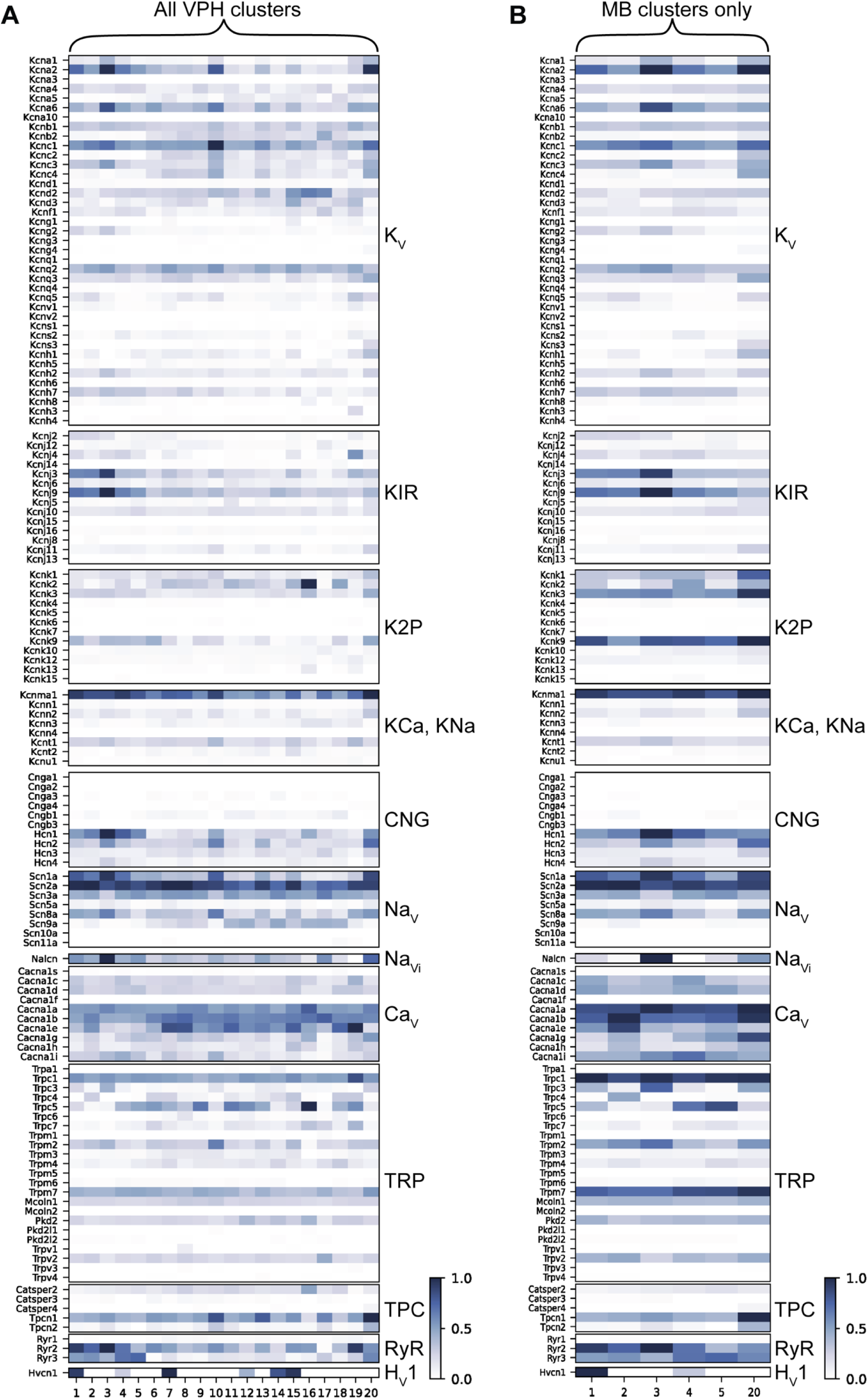
Transcripts for ion channel family members are differentially expressed across the VPH. **A)** Heat map showing the mean expression of all ion channels across the VPH, scaled to the maximum value within each channel family. **B)** Heat map showing the mean expression of ion channels within the clusters representing only the MB, scaled independently of A.

Generally, transcripts for Na_V_ channels are expressed throughout the VPH with the exception of Na_V_1.8 (*Scn10a*), Na_V_1.9 (*Scn11a*), and Na_V_1.5 (*Scn5a*), which are either absent or undetectable (Fig. 3A). Na_V_1.2 (*Scn2a*) is highly expressed across the VPH, consistent with its widespread expression throughout the brain (Shin et al., 2019). Na_V_1.3 (*Scn3a*) is moderately expressed in the VPH, with transcripts detected in most subregions. Other Na_V_ channel transcripts exhibited more cluster-enriched expression patterns across the VPH. The expression pattern of Na_V_1.1 (*Scn1a*) is particularly interesting as, while transcripts do appear elsewhere in the VPH, it is most strongly expressed in the 6 clusters representing the MB. This is a unique pattern, as this channel is mostly found in inhibitory neurons in the hippocampus and cortex (Yamagata et al., 2017), but in our dataset, appears to be enriched in the exclusively excitatory neurons of the MB. Na_V_1.7 (*Scn9a*) transcripts, in contrast, appear mostly in regions of the VPH outside of the MB with little to no expression detected in the MB clusters.

Transcripts for both Ca_V_ and ryanodine receptors show expression both throughout the VPH and, in the case of the latter, specifically within MB clusters. The Ca_V_ channel transcripts *Cacna1a* and *Cacna1b* are detected at moderate levels across the VPH, while *Cacna1e* shows a pattern of enrichment outside of the MB clusters (Fig. 3A). Notably, transcripts for the low-threshold, T-type calcium channel *Cacna1g* are detected at lower levels in the VPH but appear to be enriched within the MB LM cluster. These Ca_V_ transcripts identified in our sequencing data represent one potential explanation for the burst firing previously described in some MB neurons (Alonso and Llinás, 1992; Llinás and Alonso, 1992). Of the ryanodine receptors, transcripts for *Ryr2* are seen throughout the VPH while *Ryr3* is specifically enriched within the MB clusters. RyR3 has an established role in synaptic plasticity, learning and memory, and social behaviors (Balschun et al., 1999; Galeotti et al., 2008; Matsuo et al., 2009).

Most channel transcripts belonging to the CNG family (*Cnga* and *Cngb*) are not highly expressed across the VPH; however, the hyperpolarization-activated cyclic nucleotide-gated (HCN) channel, particularly *Hcn1* and *Hcn2,* show a clear pattern of enrichment in MB clusters relative to the rest of the VPH (Fig. 3A). The expression of these channels may provide an explanation for the firing patterns previously reported in MM and LM neurons (Llinás and Alonso, 1992; Alonso and Llinás, 1992) as they are often associated with burst firing patterns in the neurons in which they are expressed (Biel et al., 2009). Transcripts for HCN1 (*Hcn1*) are highly expressed in the MB clusters, with the highest expression in cluster 3. *Hcn2* is more broadly expressed across the VPH, with higher expression in the LM and dorsal MnM clusters. The remaining two channels appear at much lower levels, with *Hcn3* being almost absent from the MB clusters and *Hcn4* not very highly expressed across the VPH.

Several members of these ion channel families are encoded by genes implicated in NDDs (Simkin et al., 2022). Notably, transcripts for several of the genes most frequently associated with NDDs, including *Scn1a, Kcnq2*, and *Hcn1*, are highly expressed or enriched within specific MB subpopulations. Although the phenotypes associated with these disorders are generally attributed to dysfunction of cortical and hippocampal circuits, the established role of the MB in memory function raises the possibility that altered excitability within MB neurons may also influence memory-related phenotypes. To better define the molecular organization of these candidate neuronal populations, we next examined the anatomical distribution of these ion channel transcripts and related family members using FISH, providing spatial context for the expression patterns identified by scRNAseq.

### NDD-associated Na_V_ channels show unique patterns of expression along the rostrocaudal axis of the MB

Given that the scRNAseq data showed patterns of differential expression for multiple Na_V_ channels, we used FISH to further investigate transcript expression for individual channels in five representative coronal sections across the rostrocaudal extent of the MB **(Fig. 4A)**. As mentioned, *Scn1a* stood out as uniquely expressed in the MB, as this region is composed entirely of glutamatergic neurons in mice and Na_V_1.1 has been shown to be enriched in inhibitory neurons based on hippocampal and cortical data (Wang et al., 2017; Mantegazza et al., 2021). *Scn1a* transcripts were found in all MB clusters but were enriched within LM and dorsal MnM subpopulations relative to the others **(Fig. 4Bi)**. FISH revealed that, while this was the case in more anterior MB sections **(Fig. 4Bii-iv)**, *Scn1a* transcripts appeared to be abundant across all subregions in more posterior sections **(Fig. 4Bv-vi)**. Less surprising was the expression of *Scn2a*, typically expressed in excitatory neurons (Mantegazza et al., 2021), which was highly expressed across all MB clusters. *Scn2a* signal is also consistently high across all subregions from anterior to posterior **(Fig. 4Ci-vi)**. *Scn8a*, expressed in both glutamatergic and GABAergic neurons in other brain regions (Mantegazza et al., 2021), is found in multiple MB clusters and, similar to *Scn1a*, appears to be enriched in LM and dorsal MnM subpopulations. While *Scn8a* signal appears to be slightly higher in the LM of anterior sections **(Fig. 4Dii-iii)**, FISH showed relatively even expression across all MB subdivisions.

**Figure 4:**
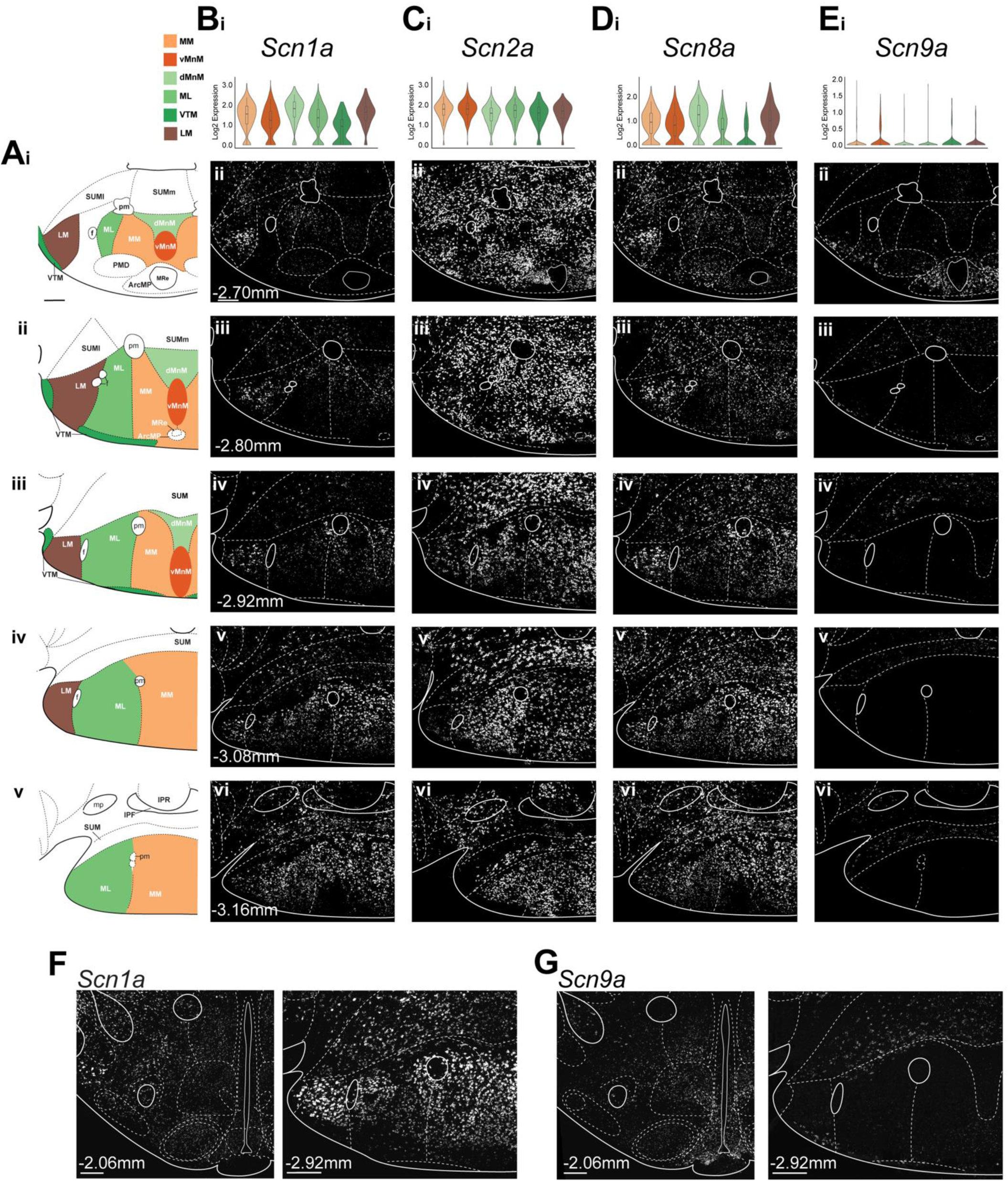
Transcripts for NaV channels are differentially expressed throughout the MB. **A)** Representative anatomical maps of the MB from anterior (i) to posterior (v) (scale bar=200 μm) **B)** Violin plot representing the scRNAseq data for *Scn1a* in each MB cluster showing the distribution of log-normalized expression. Box plots indicate the median, interquartile range, and 5^th^-95^th^ (i) FISH showing the expression of *Scn1a* transcripts through the volume of the MB (ii-vi) (n=3 mice) (scale bar=200 μm) **C)** Violin plot of *Scn2a* expression in MB clusters (i) and representative images showing its expression within the MB (ii-vi) (n=5 mice). **D)** Violin plot of *Scn8a* in the MB (i) along with FISH images of its expression throughout the MB (ii-vi) (n=3 mice). **E)** Violin plot showing little to no detectable expression of *Scn9a* transcripts in the MB (i), is confirmed by images from FISH for this transcript (ii-vi) (n=3 mice). **F)** FISH images comparing *Scn1a* expression in the MB (right) to its expression in more anterior portions of the hypothalamus (left) (scale bars=200 μm). **G)** Representative FISH images showing *Scn9a* expression in the MB (right) relative to its expression in an anterior region of the hypothalamus (left) (scale bars=200 μm).

In contrast to the other Na_V_ channels examined, *Scn9a* showed little to no expression in MB clusters **(Fig. 4Ei)**, despite being expressed elsewhere in the VPH (Fig. 3A). In the anterior-most section, *Scn9a* signal is enriched in the arcuate nucleus, consistent with previous findings showing its expression in this region (Branco et al, 2016). Conversely, within the same anterior sections, there is sparse signal in the MB **(Fig. 4Eii)**. In more posterior sections, signal is visible in the supramammillary nucleus but not in the MB **(Fig. 4Eiii-vi)**.

To directly compare Na_V_ channel transcripts with contrasting patterns of MB enrichment, we examined *Scn1a*, which is enriched in the MB, and *Scn9a*, which is very sparse, across sections containing the MB and more rostral hypothalamic regions in the same mouse. Direct comparison of *Scn1a* expression in the MB and more rostral hypothalamic regions revealed an apparent enrichment of *Scn1a* within the MB **(Fig. 4F)**. Conversely, *Scn9a* showed the opposite distribution, with prominent expression in medial regions of the rostral hypothalamus, as well as in the supramammillary nucleus, but little to no expression within the MB **(Fig. 4G)**. Taken together, these data demonstrate distinct patterns of Na_V_ channel transcript expression within the MB, with several showing relative enrichment in the MB compared with other hypothalamic regions. Of particular interest among these channels is the unique expression pattern of *Scn1a* and *Scn2a*. Specifically, we asked whether MB neurons co-express these channels or if they are simply expressed by separate subpopulations within the same subregions.

### *Scn1a* and *Scn2a* are co-expressed across several MB subdivisions

Given the broad expression of both *Scn1a* and *Scn2a* across MB subregions, we next asked how their expression was distributed among molecularly defined MB neuronal populations. To this end, we carried out multiplex FISH probing for transcriptomic markers of MB subregions, *Tac2* for LM, *Pvalb* for LM and MnM, *Calb1* for ML, and *Nts* for MM, along with either *Scn1a* or *Scn2a*. Representative images of a central MB section show that *Scn1a* is expressed in a large proportion of each population within each subregion of the MB **(Fig. 5A)**, consistent with our transcriptomic data. Within the LM, defined by neurons expressing *Pvalb, Tac2* or both, *Scn1a* is expressed in 88% of neurons. In the neighboring ML, 73% of *Calb1*-expressing neurons also express *Scn1a,* and 85% of MM *Nts* neurons express *Scn1a*. MnM *Pvalb*-expressing neurons showed the highest co-expression of *Scn1a,* with 93% of these neurons being co-labeled for *Scn1a* transcripts. As expected, we found that *Scn2a* is expressed in all the assessed cell clusters and subregions **(Fig. 5B)**. We found that 78% of the LM neurons assessed also expressed *Scn2a* while 76% ML *Calb1* neurons express *Scn2a*. *Scn2a* is found in 83% of MM *Nts*+ neurons, and the same is true of MnM *Pvalb* neurons. The high levels of expression within each cell type and subregion also suggest that these neurons may also be co-expressing *Scn1a* and *Scn2a,* a trait not often seen in the hippocampus and cortex (Yamagata et al, 2017). Together, these channels may both play a key role in action potential firing in MB neurons and in the integration and propagation of signal between MB neurons and their target structures. Having characterized Na_V_ channel expression, we next examined voltage-gated potassium (K_V_) channels, which further shape action potential firing and regulate neuronal excitability.

**Figure 5:**
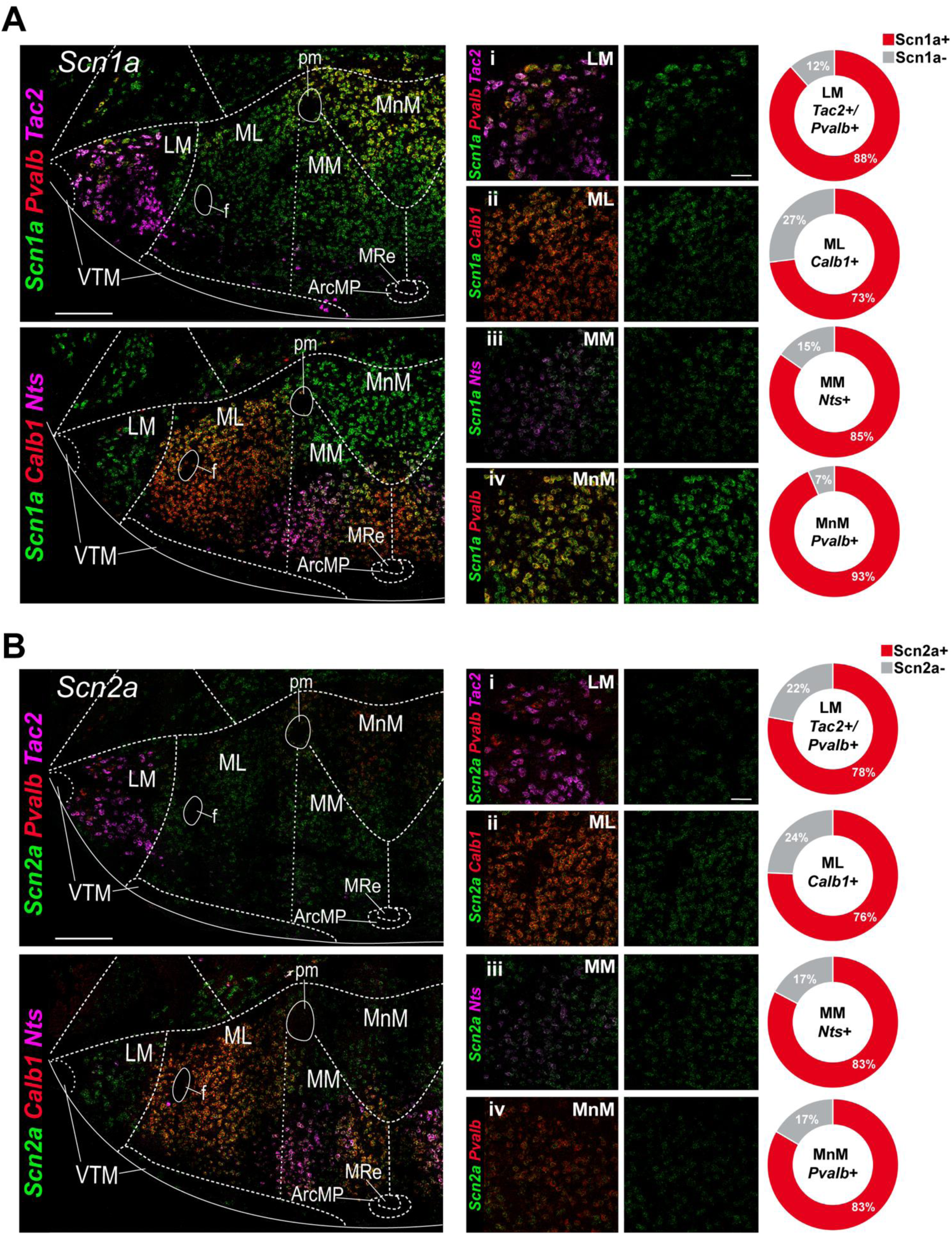
*Scn1a* and *Scn2a* are highly expressed within subpopulations of MB neurons. **A)** 40x confocal image of a section co-stained for *Scn1a* (green), *Pvalb* (red) and *Tac2* (magenta) [top left] with clear co-labelling for both *Pvalb* and *Scn1a* transcripts within the MnM indicated by the yellow signal. A second 40x image taken of a section from the same animal co-stained with *Scn1a*, *Calb1* (red), and *Nts* (magenta) [bottom left] shows co-labeling of *Scn1a* with *Calb1* in more dorsal portions of the ML (yellow signal) as well as *Scn1a* with *Nts* in the MM (white signal) (scale bars=200 µm). 100x images in the LM (i), ML (ii), MM (iii) and dorsal MnM (iv) showing co-labeling with cell type markers for each subregion as well as the expression pattern of *Scn1a* alone within each subregion [middle panel] and quantification of expression within each cell type [far right] (n=5 mice, 283 cells [LM], 610 cells [MnM], 467 cells [ML], 445 cells [MM]) (scale bar=50 µm) **B)** 40x image of a section co-stained for *Scn2a* (green), *Pvalb* (red) and *Tac2* (magenta) [top left]. A second 40x image taken of a section from the same animal co-stained with *Scn2a*, *Calb1* (red), and *Nts* (magenta) [bottom left] co-labeling is visible with *Calb1* in the ML (yellow signal) and *Nts* in the MM (white signal). 100x images in the LM (i), ML (ii), MM (iii) and dorsal MnM (iv) showing co-labeling with cell type markers for each subregion as well as the expression pattern of *Scn2a* alone within each subregion [middle panel] and quantification of expression within each cell type [far right] (n=5 mice, 304 cells [LM], 699 cells [MnM], 522 cells [ML], 560 cells [MM])

### NDD-associated K_V_ channels are differentially expressed within the MB

Transcripts encoding K_V_ channels also exhibited distinct patterns of expression across MB subpopulations **(Fig. 6A)**. To further characterize these patterns, we focused on three K_V_ channel genes implicated in NDDs: *Kcna2*, *Kcnq2*, and *Kcnq3*. Among these channels, both *Kcna2* and *Kcnq2* were found in each MB cluster, with the former showing higher expression in the dorsal MnM and LM and the latter showing slight enrichment in the dorsal MnM **(Fig. 6Bi and Ci)**. Conversely, *Kcnq3* is not highly expressed in the MB with the exception of the LM **(Fig. 6Di)**. FISH revealed cells expressing *Kcna2* transcripts in all MB subregions, with LM enrichment being more visible in anterior sections and showing diminishing signal in more posterior sections **(Fig. 6Bii-vi)**. A pattern of higher expression in the MnM is more apparent in anterior and middle sections, while ML expression remained consistent throughout. *Kcnq2* signal is also visible in all MB subregions, though the majority of *Kcnq2*+ neurons appear within the medial nucleus (MnM, MM and ML) with sparse expression in the LM **(Fig. 6Cii-vi)**. In contrast, *Kcnq3*-expressing cells are concentrated in the LM in more rostral sections, with comparatively sparse expression throughout the medial nucleus **(Fig. 6Dii-vi)**. K_V_7.2 and K_V_7.3 are frequently found together in other regions of the brain and form heteromeric channels (Wang et al., 1998). However, the distinct expression patterns of *Kcnq2* and *Kcnq3* raise the possibility that their relative expression, and potentially K_V_7 channel composition, differs across MB neuronal populations. We therefore used multiplex FISH to determine the extent to which *Kcnq2* and *Kcnq3* transcripts are expressed within molecularly defined neuronal populations across MB subregions.

**Figure 6:**
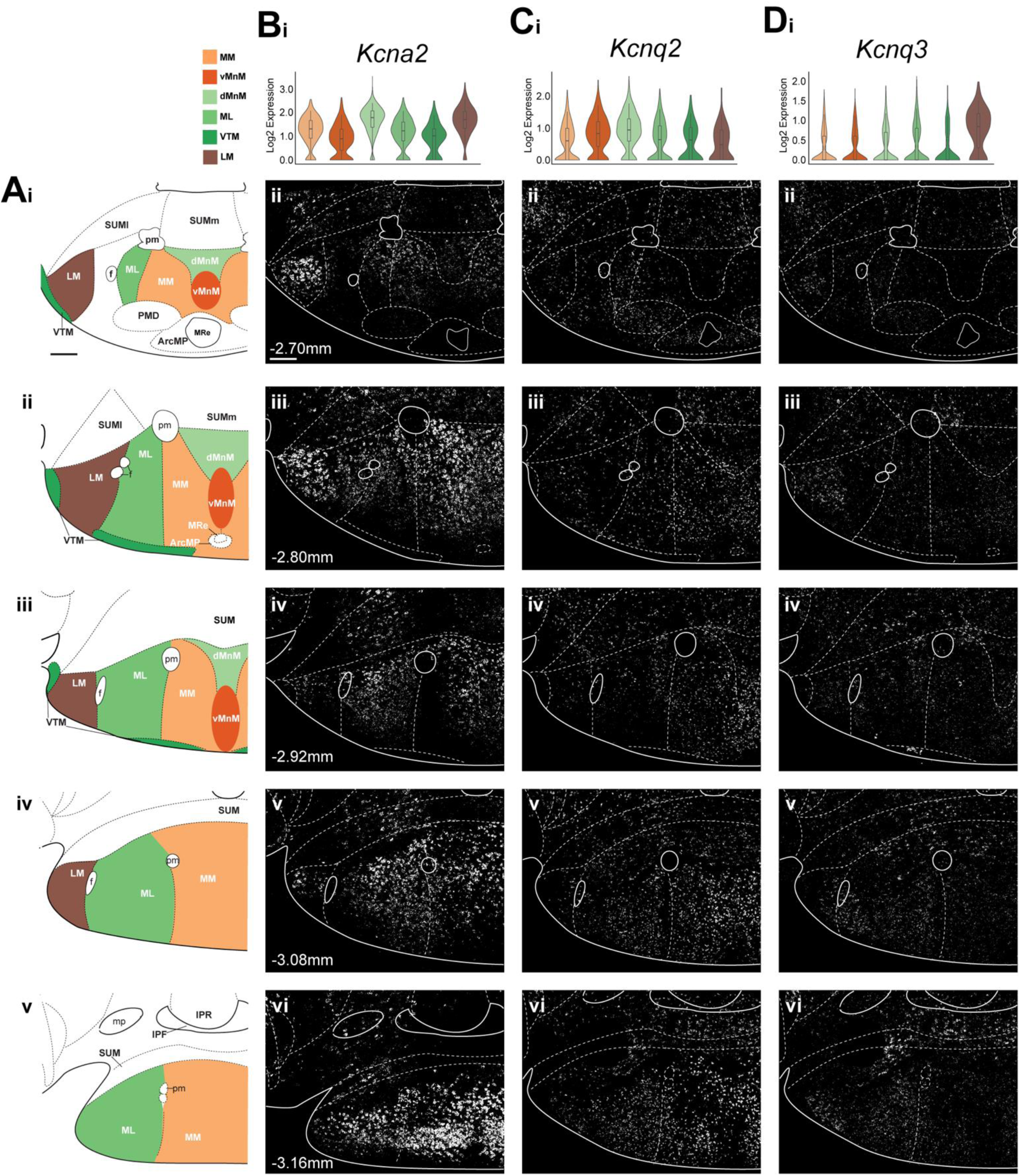
Transcripts for KV channels are differentially expressed throughout the MB. **A)** Representative anatomical maps of the MB from anterior (i) to posterior (v) (scale bar=200 μm) **B)** Violin plot showing *Kcna2* expression detected via scRNAseq (i) FISH shows strong signal within the LM in anterior and middle MB sections (ii-iv) with additional signal in the medial nucleus (both ML and MM) (v-vi) (n=3) (scale bar=200 μm). **C)** Violin plots of *Kcnq2* expression (i) with FISH showing it is evenly distributed throughout subregions of the MB showing slightly higher expression in the MnM and diminished expression in the LM (ii-vi) (n=3 mice). **D)** Violin plot showing *Kcnq3* RNA in the MB (i) with FISH probing for *Kcnq3* transcripts showing LM enrichment in anterior sections (ii-iii), with some expression in other subregions towards the posterior end of the MB (v-vi) (n=3 mice).

### *Kcnq2* is highly expressed across the MB while *Kcnq3* expression is sparse

The differential expression of *Kcnq2* and *Kcnq3* suggests that subpopulations of MB neurons may express one or the other. Evidence for this type of expression has only been observed in select populations of neurons throughout the brain (Schwarz et al., 2006; King et al., 2014; Sun et al., 2019; Goff and Goldberg, 2019). Upon further inspection of the neuronal subpopulations within which these channels are expressed, we found that the majority of MB neurons express the former, with fewer expressing the latter **(Fig. 7A and B)**. Of note, *Kcnq2* is enriched in MnM *Pvalb*+ neurons (91%) **(Fig. 7Aiv)**, consistent with the pattern seen in single-plex FISH and the single-cell data, which showed it as more strongly expressed in the dorsal MnM population. Conversely, comparatively few (29%) *Pvalb*+ neurons in the MnM express *Kcnq3* **(Fig. 7Biv)**. This differential expression raises the possibility that K_V_7.2-containing channels in many MnM *Pvalb*+ neurons may function independently of K_V_7.3, although channel subunit composition cannot be determined from transcript expression alone. In the LM, most of the assessed neurons also express *Kcnq2* (81%) **(Fig. 7Ai)** while very few of these neurons (45%) also express *Kcnq3* **(Fig. 7Bi)**. The LM thus displays a similar pattern to that of the dorsal MnM neurons. In the medial nucleus, 65% of *Calb1-expressing* neurons are *Kcnq2*+ **(Fig. 7Aii)**, while only 32% are *Kcnq3*+ **(Fig. 7Bii)**. Furthermore, we found that 76% of *Nts*+ neurons are also positive for *Kcnq2* **(Fig. 7Aiii)** with only 22% expressing *Kcnq3* **(Fig. 7Biii)**. Overall, our FISH data demonstrate that *Kcnq2* is broadly expressed across MB neuronal populations, whereas *Kcnq3* is expressed in a smaller proportion of neurons and shows greater enrichment within the LM. These distinct expression patterns suggest that the relative contribution of K_V_7.2 and K_V_7.3 to K_V_7 channel composition may differ across MB neuronal populations.

**Figure 7:**
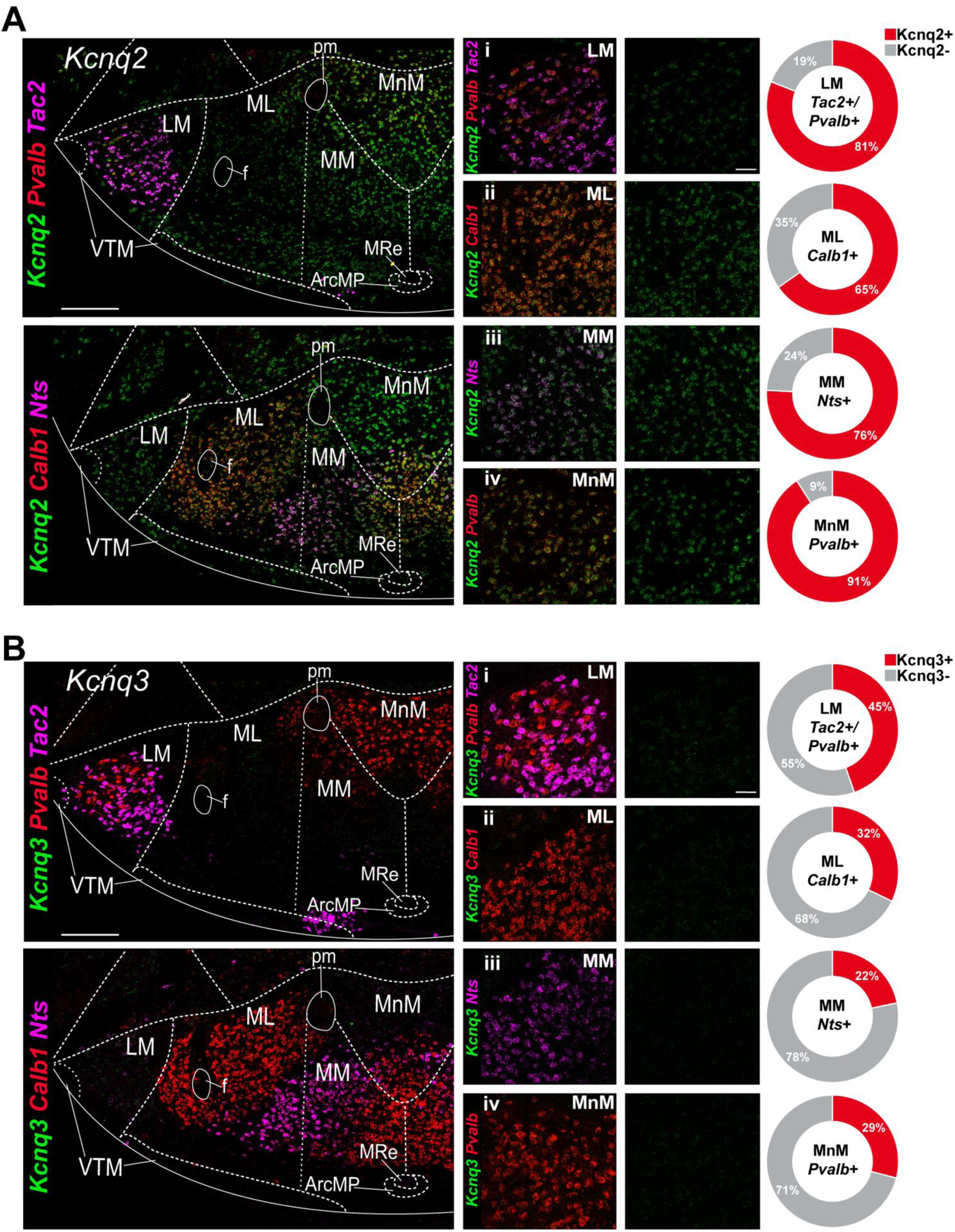
*Kcnq2* but not *Kcnq3* is expressed within multiple MB neuronal subpopulations. **A)** 40x image of a section co-stained for *Kcnq2* (green), *Pvalb* (red) and *Tac2* (magenta) [top left] co-labelling for both *Pvalb* and *Kcnq2* transcripts is visible within dorsal portions of MnM (yellow signal). A second 40x image taken of a section from the same animal co-stained with *Kcnq2*, *Calb1* (red), and *Nts* (magenta) [bottom left] shows co-labeling of *Kcnq2* with *Calb1* throughout the ML (yellow signal) as well as *Kcnq2* with *Nts* in the MM (white signal) (scale bars=200 µm). 100x images in the LM (i), ML (ii), MM (iii) and dorsal MnM (iv) showing co-labeling with cell type-specific transcripts for each subregion as well as the expression pattern of *Kcnq2* alone within each subregion [middle panel] and quantification of expression within each cell type [far right] (n=5 mice, 322 cells [LM], 692 cells [MnM], 682 cells [ML], 631 cells [MM]) (scale bar=50 µm). **B)** 40x image of a section co-stained for *Kcnq3* (green), *Pvalb* (red) and *Tac2* (magenta) [top left]. A second 40x image taken of a section from the same animal co-stained with *Kcnq3*, *Calb1* (red), and *Nts* (magenta) [bottom left] Both images show very little expression of *Kcnq3* transcripts. 100x images in the LM (i), ML (ii), MM (iii) and dorsal MnM (iv) showing co-labeling with cell type markers for each subregion as well as the expression pattern of *Kcnq3* alone within each subregion [middle panel] and quantification of expression within each cell type [far right] (n=5 mice, 292 cells [LM], 1156 cells [MnM], 659 cells [ML], 731 cells [MM]).

### HCN channel transcripts are enriched in MB subdivisions

The final NDD-associated ion channel family that we investigated was the HCN channel family **(Fig. 8A)**. These channels are often associated with rhythmic bursting and produce a hyperpolarization-activated current known as I_h_ (Biel et al., 2009). Previous electrophysiological data showing rhythmic bursts and rebound bursts in subpopulations of MB neurons (Alonso and Llinás, 1992) raise the possibility that these channels may be involved in determining their intrinsic firing properties. Single-cell data for all HCN channels shows that transcripts for *Hcn1* can be found in all MB subdivisions, with slightly stronger expression in dorsal MnM neurons **(Fig. 8Bi)**, while *Hcn2* is most highly expressed in LM neurons and found at lower levels elsewhere **(Fig. 8Ci)**. *Hcn3* appears at low levels in most MB clusters with relatively higher expression in LM neurons **(Fig. 8Di)**. *Hcn4* also shows a pattern of generally low expression within MB clusters, with slightly higher expression in dorsal MnM neurons **(Fig. 8Ei)**. FISH for *Hcn1* shows expression in all MB subregions. There is a pattern of *Hcn1* enrichment in the LM and dorsal MnM of more anterior sections, while more posterior sections show even expression across subregions **(Fig. 8Bii-vi)**. Analysis of *Hcn2* reveals an unexpected pattern of expression: while there is a high density of signal in the LM, there is clearly very strong signal throughout the medial nucleus as well **(Fig. 8Cii-vi)**, in contrast to the neuronal single-cell data; however, we found that several non-neuronal populations, specifically oligodendrocytes, show enriched expression of *Hcn2* and may be the reason for the additional signal visible in the medial MB. For *Hcn3,* the anterior-most and posterior-most sections show the expected sparse expression; however, the central sections show higher expression throughout the MB **(Fig. 8Dii-vi)**. Finally, *Hcn4* shows very little expression within MB subregions except for the MnM, which shows sparse signal **(Fig. 8Eii-iv)** that becomes more pronounced in the more posterior sections within the ML and MM **(Fig. 8Ev-vi)**.

**Figure 8:**
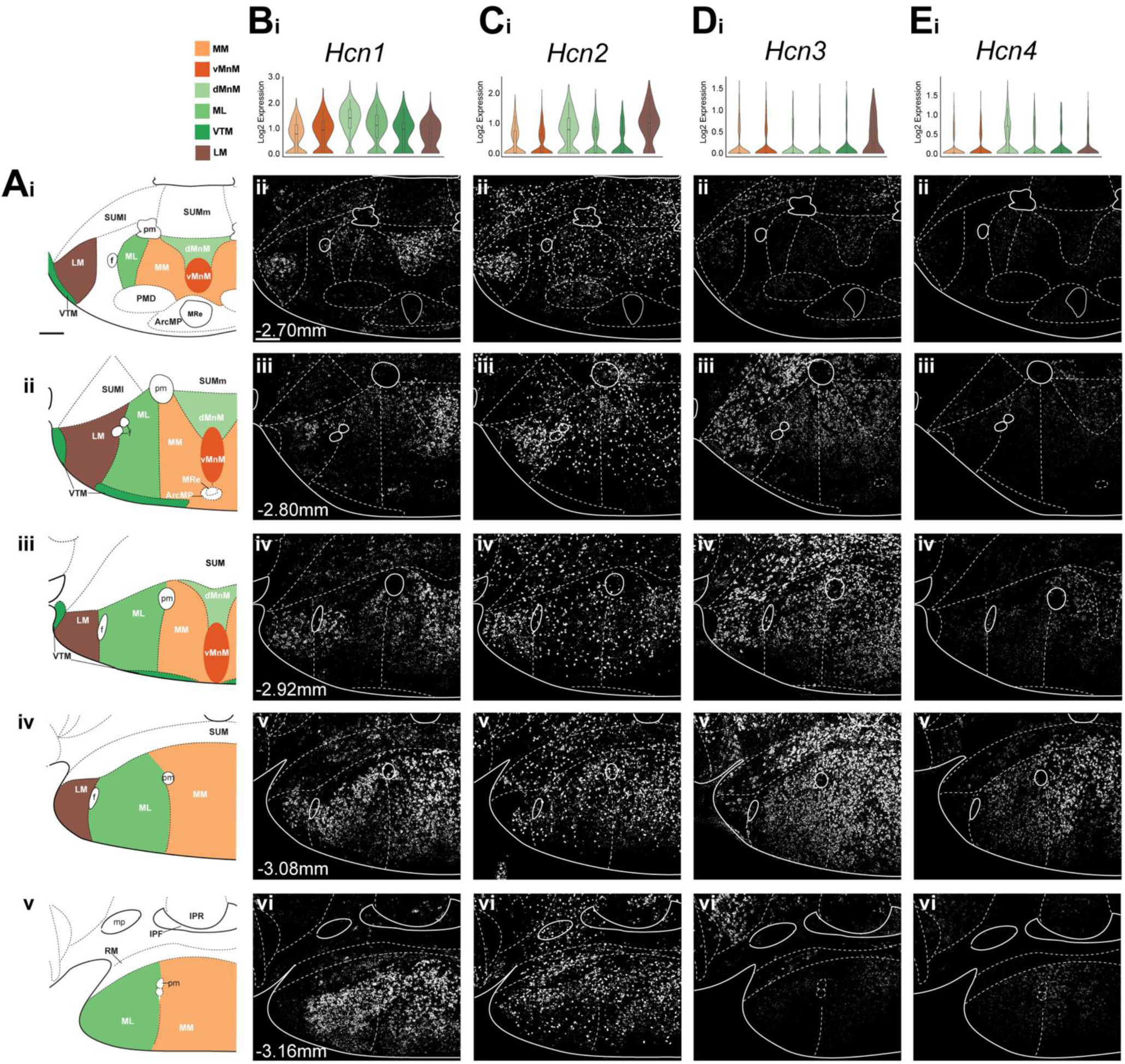
Differential expression of transcripts for HCN channels throughout the MB. **A)** Representative anatomical maps of the MB from anterior (i) to posterior (v) (scale bar=200 μm) **B)** Violin plot of *Hcn1* expression in the MB (i) FISH shows it is enriched in the MnM and the LM a pattern that is visible in anterior and middle (ii-iv) MB sections with greater signal becoming visible throughout the medial nucleus in posterior (v-vi) sections (n=6 mice) (scale bar=200 μm). **C)** Violin plots showing *Hcn2* RNA in the MB (i) with FISH showing its expression throughout the MB (ii-vi) revealing that while there is a concentration of signal in the LM, there is additional signal visible throughout the medial nucleus (n=3 mice). **D)** violin plots for *Hcn3* transcripts (i) along with representative FISH images showing its expression in the MB (ii-vi) (n=4 mice). **E)** Violin plots show very low expression of *Hcn4* in the MB (i) FISH shows some signal in the MnM of anterior and middle sections (ii-iv) with a slight increase going more posterior (v) but then a return to low signal in the posterior-most section (vi) (n=3 mice).

### *Hcn1* exhibits enrichment in MB *Pvalb* neurons

Given that *Hcn1* showed the strongest and most distinct pattern of expression among HCN family members, we next examined its distribution across molecularly defined MB neuronal populations using multiplex FISH **(Fig. 9)**. In general, most of the populations we stained for co-expressed *Hcn1.* Still, a clear pattern of enrichment in *Pvalb*+ MnM neurons was revealed, with 76% of these neurons expressing *Hcn1* **(Fig. 9Aiv)** while only 49% of LM neurons were *Hcn1*+ **(Fig. 9Ai)**. Consistent with both the single-cell and single-plex FISH data, 61% of *Calb1*-expressing neurons in the ML express *Hcn1,* while only 44% of *Nts*+ neurons are positive for *Hcn1*. Together, these data show that *Hcn1* is expressed across multiple MB neuronal populations but is more frequently detected in *Pvalb*-expressing neurons.

**Figure 9:**
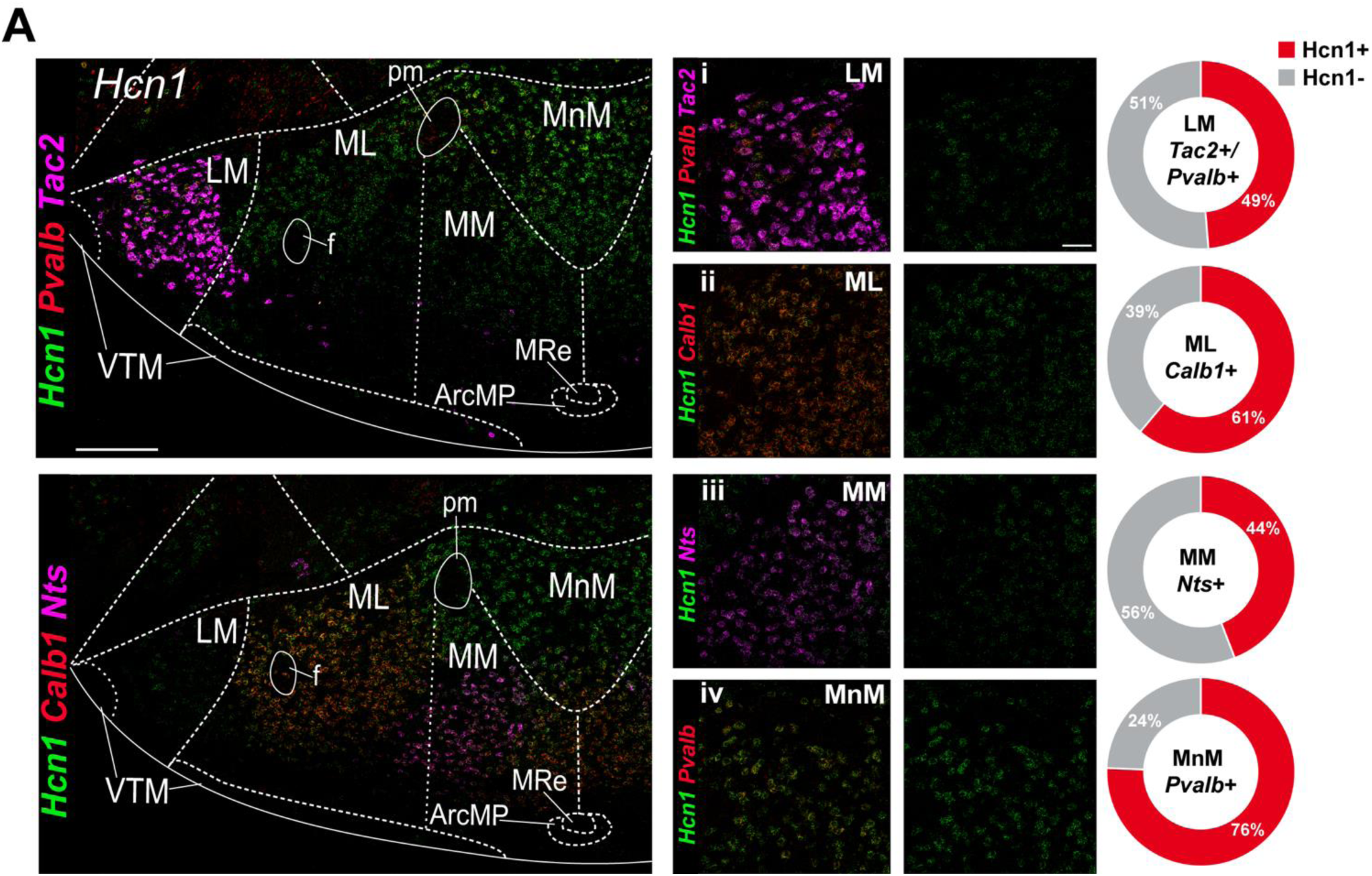
*Hcn1* shows a pattern of enrichment in MB *Pvalb+* neurons. **A)** 40x image of a section co-stained for *Hcn1* (green), *Pvalb* (red) and *Tac2* (magenta) [top left] some co-labelling for *Pvalb* and *Hcn1* transcripts in dorsal portions of the MnM indicated by the yellow signal. A second 40x image taken of a section from the same animal co-stained with *Hcn1*, *Calb1* (red), and *Nts* (magenta) [bottom left] with co-labeling of Hcn1 with *Calb1* throughout the ML (yellow signal) (scale bars=200 µm). 100x images in the LM (i), ML (ii), MM (iii) and dorsal MnM (iv) showing co-labeling with cell type-specific markers for each subregion as well as the expression pattern of *Hcn1* alone within each subregion [middle panel] and quantification of expression within each cell type [far right] (n=5 mice, 271 cells [LM], 854 cells [MnM], 743 cells [ML], 743 cells [MM]) (scale bar=50 µm).

### NaV, KV and HCN channel transcripts in LM *Pvalb*+ and *Tac2*+ subpopulations

Our initial analyses characterized ion channel transcript expression across the LM as a whole. Given that the LM contains molecularly distinct *Pvalb*+ (enriched in the dorsal LM) and *Tac2*+ (enriched in the more central/ventral LM) neuronal populations, we next asked whether these populations exhibit differential patterns of expression. When split into populations based on the expression of *Pvalb* or *Tac2* alone, we found that more *Pvalb*+ neurons express *Scn1a* (88%) than *Tac2*+ neurons (69%) **(Fig. 10A)**. For *Scn2a*, we found that a more similar proportion of *Pvalb*+ and *Tac2*+ LM neurons express Scn2a (72% and 76%, respectively) **(Fig. 10B)**. For *Kcnq2*, a similar proportion of *Pvalb*+ and *Tac2*+ neurons were *Kcnq2*+ (78% and 72% respectively **(Fig. 10C)**. Although *Kcnq3* was also similar between the two subpopulations (53% and 42% respectively), the proportion was substantially less than *Kcnq2* **(Fig. 10D)**. Finally, we found that a greater proportion of LM *Pvalb*+ neurons express *Hcn1* (62%) than LM *Tac2* neurons (34%) **(Fig. 10E)**. Together with our data from Fig. 9, these results suggest an enrichment of *Hcn1* across dorsally localized LM and MnM *Pvalb*+ neurons.

**Figure 10:**
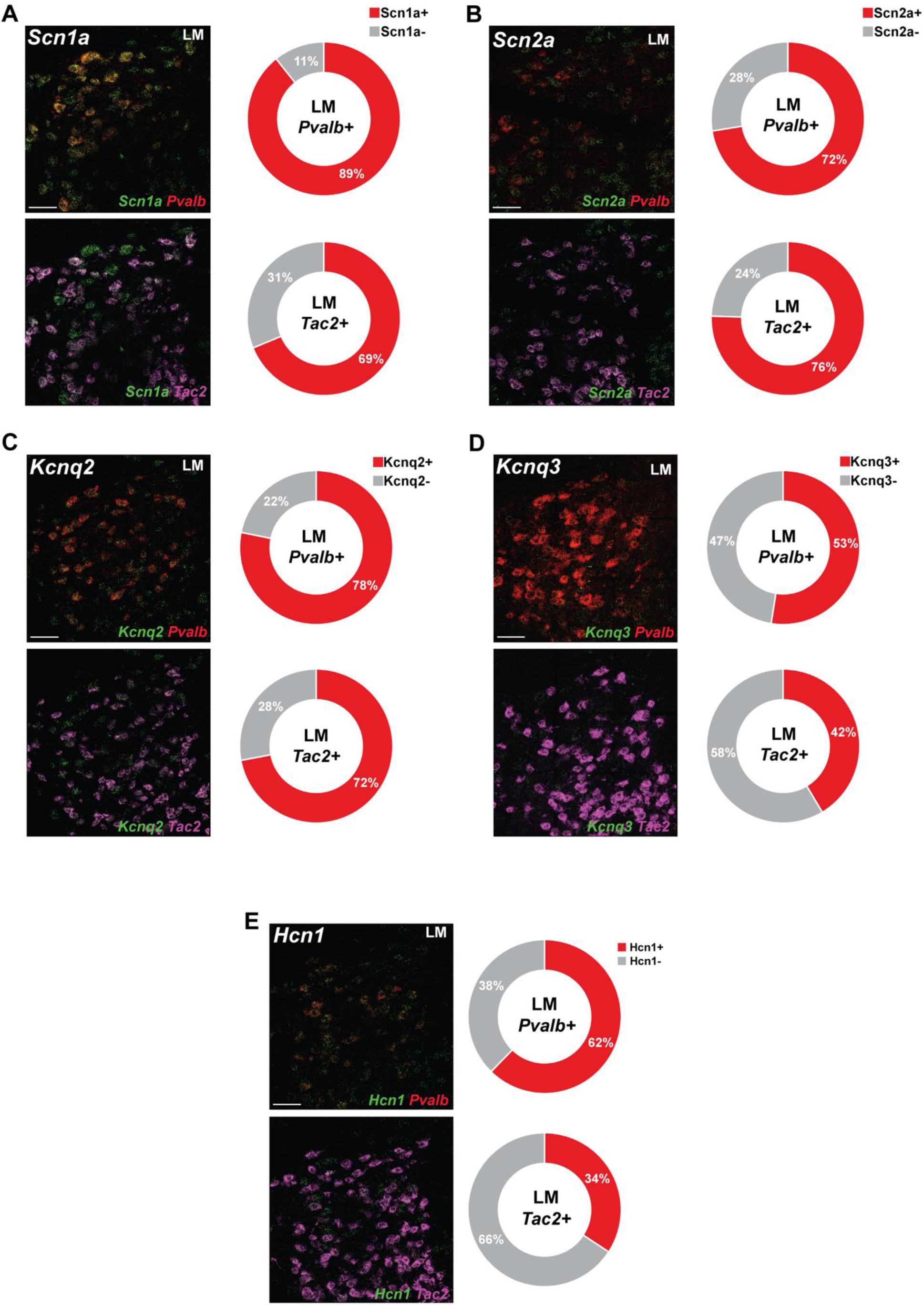
NaV, KV and HCN channel transcripts in LM *Pvalb*+ and *Tac2*+ subpopulations. **A)** Left, confocal images (100x) of multiplex FISH in the LM staining for *Scn1a* (green), *Pvalb* (red), and *Tac2* (magenta). Top panel shows overlay of *Scn1a* and *Pvalb* signal and bottom panel shows overlay of *Scn1a* and *Tac2* [scale bar=50 µm]. Accompanying quantification for neurons only expressing *Pvalb* (top right; n=5 mice 168 cells) and neurons only expressing *Tac2* (bottom right; n=5 mice; 48 cells) **B)** 100x confocal images of multiplex FISH in the LM staining for *Scn2a* (green), *Pvalb* (red), and *Tac2* (magenta). Top panel shows overlay of *Scn2a* and *Pvalb* signal and bottom panel shows overlay of *Scn2a* and *Tac2* [scale bar=50 µm] with accompanying quantification for *Pvalb* only neurons (top right; n=5 mice; 218 cells) and *Tac2* only (bottom right; n=5 mice; 49 cells) **C)** Confocal images (100x) of multiplex FISH in the LM staining for *Kcnq2* (green), *Pvalb* (red), and *Tac2* (magenta). Top panel shows overlay of *Scn1a* and *Pvalb* signal and bottom panel shows overlay of *Kcnq2* and *Tac2* [scale bar=50 µm]. Accompanying quantification for neurons only expressing *Pvalb* (top right; n=5 mice; 174 cells) and neurons only expressing *Tac2* (bottom right; n=5 mice; 50 cells) **D)** 100x confocal images of multiplex FISH in the LM staining for *Kcnq3* (green), *Pvalb* (red), and *Tac2* (magenta). Top panel shows overlay of *Scn2a* and *Pvalb* signal and bottom panel shows overlay of *Kcnq3* and *Tac2* [scale bar=50 µm] with accompanying quantification for *Pvalb* only neurons (top right; n=5 mice; 160 cells) and *Tac2* only (bottom right; n=5 mice; 41 cells) **E)** Confocal images (100x) of multiplex FISH in the LM staining for *Hcn1* (green), *Pvalb* (red), and *Tac2* (magenta). Top panel shows overlay of *Hcn1* and *Pvalb* signal and bottom panel shows overlay of *Hcn1* and *Tac2* [scale bar=50 µm]. Accompanying quantification for neurons only expressing *Pvalb* (top right; n=5 mice; 175 cells) and neurons only expressing *Tac2* (bottom right; n=5 mice; 38 cells).

### LM and MnM neurons exhibit unique intrinsic properties

The differential expression of voltage-gated ion channel transcripts across MB neuronal populations led us to examine whether neurons in distinct MB subregions also differ in their intrinsic electrophysiological properties. To this end, we performed whole-cell current-clamp recordings from neurons in the dMnM and LM **(Fig. 11A)**, two regions containing *Pvalb*+ neuronal populations but exhibiting distinct ion channel expression profiles. Applying current steps, we found that LM neurons were generally less excitable than MnM neurons. Typically, LM neurons fired several action potentials near the beginning of the step before falling silent immediately after **(Fig. 11Ai)**. In contrast, MnM neurons were found to initially burst in response to current injections but quickly enter a pattern of rapid, tonic firing **(Fig. 11Aii)**. Consistent with their lower excitability, LM neurons fired significantly fewer action potentials than MnM neurons across equivalent current injections, as reflected in the input-output relationship **(Fig. 11B)**. For example, at 95 pA, LM neurons fired 7.82 ± 3.82 action potentials (n = 17), compared with 37.8 ± 4.89 in MnM neurons (n = 30) (mean ± SEM; p < 0.0001). Given that our single-cell data showed enrichment of transcripts for the leak potassium channels *Kcnk1*, *Kcnk3*, and *Kcnk9*, we wanted to determine whether these characteristics were related to differences in input resistance (R_in_). Our analysis revealed that there was no difference in R_in_ between LM neurons (297 mΩ ± 47.6 with a -20 pA injection [n=16]) and MnM neurons (322 mΩ ± 28.3 with a -20 pA injection [n=29]) (p>0.9999) **(Fig. 11C)**. In line with the initially observed characteristics, rheobase for MnM vs LM neurons was 33.5 pA ± 3.35 (n=29) and 60.4 pA ± 7.54 (n=13) respectively **(Fig. 11E)**. In observing single action potential properties, we found that, along with the differences in rheobase and input-output curve, MnM neurons had similar action potential amplitudes to LM neurons (MnM=33.9 mV ± 1.18 [n=32] and LM=30.1 mV ± 1.70 [n=13]) **(Fig. 11F)** but LM neurons had significantly longer half-widths (0.724 ms ± 0.139 [n=13]) than MnM neurons (0.272 ms ± 0.014 [n=32]) (p=0.0016) **(Fig. 11G)**. Consistent with this finding, both the rise and decay times for LM neurons (rise=0.245 ms ± 0.040 [n=13]; decay=0.466 ms ± 0.097 [n=13]) were significantly higher than those of MnM neurons (rise=0.110 ms ± 0.005 [n=32]; decay=0.171 ms ± 0.009 [n=32]) (rise time p=0.0031; decay time p=0.0049) **(Fig. 11H-I)**. Fast afterhyperpolarization (fAHP) was not significantly different between the two subpopulations (LM=-43.5 mV ± 1.78 [n=13]; MnM=-40.1 mV ± 1.40 [n=32]) (p=0.2277) **(Fig. 11J)**. Together, these findings demonstrate that LM and MnM neurons, despite sharing populations of *Pvalb*+ neurons, have markedly different intrinsic electrophysiological properties. LM neurons exhibit lower excitability and broader, slower action potentials than MnM neurons. These physiological differences are consistent with the distinct ion channel expression profiles identified across these MB subregions and provide a functional basis for further investigating their underlying ionic mechanisms.

**Figure 11:**
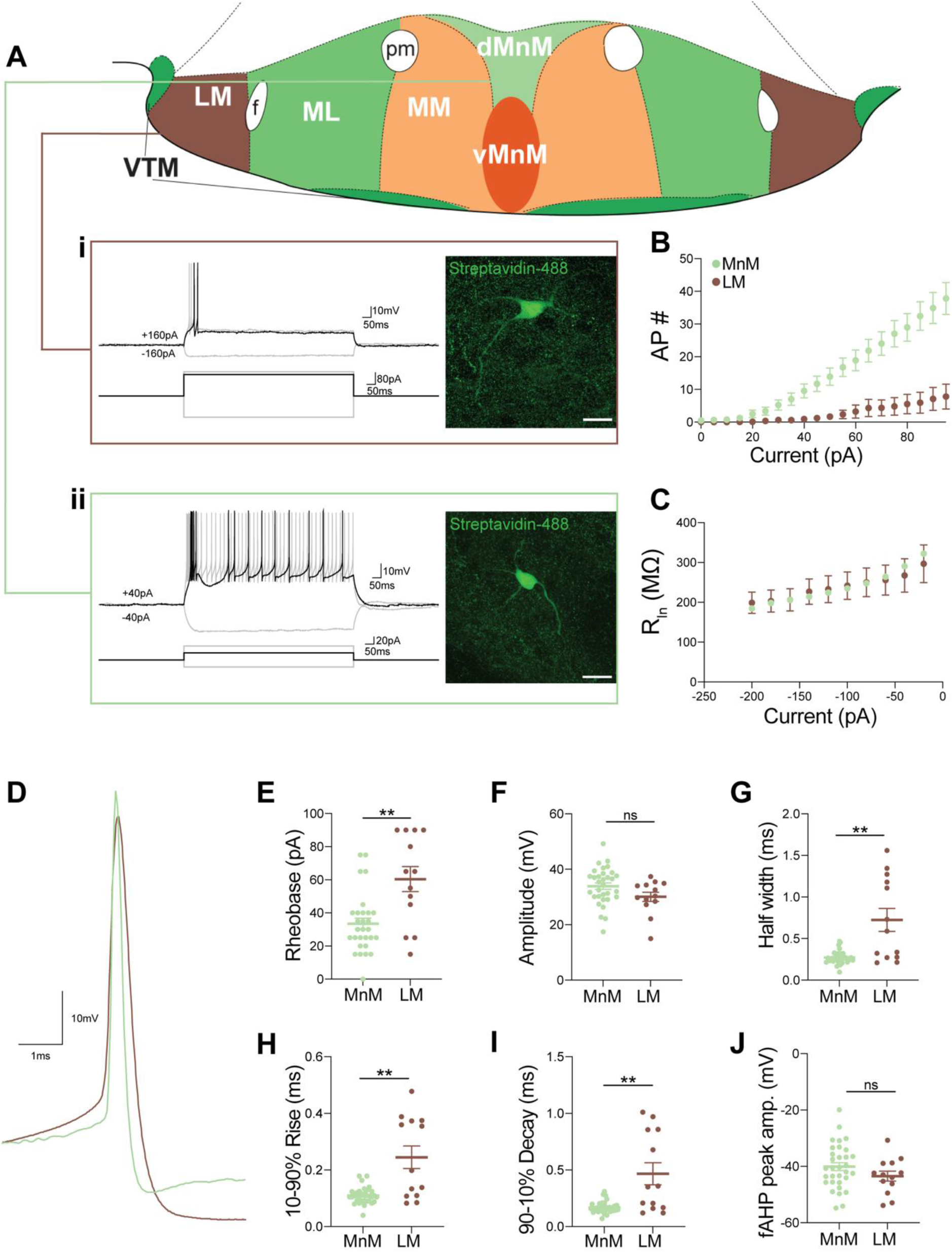
Electrophysiological characteristics of LM and MnM neurons. **A)** Representative traces from whole cell recordings of LM neurons held at -70 mV following hyperpolarizing and depolarizing steps of ±160 pA (i) and MnM neurons held at -70 mV following hyperpolarizing and depolarizing steps of ±40 pA (ii). Confocal images (100x) of recorded and biocytin-filled neurons (stained with streptavidin conjugated to Alexa-488) from each region to the right of each trace [scale bar=20 µm]. **B)** Action potential number as a function of current injected in LM neurons (n=17 cells) and MnM neurons (n=30 cells). **C)** Input resistance of recorded LM neurons (n=16 cells) and MnM neurons (n=29 cells) as a function of injected current. **D)** Overlayed single action potentials from the rheobase steps of representative LM (brown) and MnM (green) neurons. **E)** Scatterplot showing individual rheobase values for MnM (green [n=29 cells]) and LM neurons (brown [n=16 cells]). **F)** AP amplitude of all MnM neurons (n=32) and LM neurons (n=13). **G)** AP half-widths plotted for all LM neurons (n=13) and MnM neurons (n=32) **H)** Plots showing the rise times of the first APs for all LM (n=13) and MnM neurons (n=32). **I)** Decay times of the first APs fired by LM (n=13) and MnM neurons (n=32). **J)** Scatterplots showing individual fAHP peak amplitudes for MnM (n=32) and LM (n=13) neurons. In E-J, asterisks indicate statistical significance **p<0.01, ns indicates no significant difference. Statistical significance is based on non-parametric Mann-Whitney tests.

## DISCUSSION

### Summary of findings

In this study, we built upon our previous single-cell transcriptomic analysis to provide a comprehensive anatomical map of voltage-gated ion channel expression across molecularly defined MB neuronal populations using FISH. We first mapped the expression patterns of multiple cluster-enriched markers, including *Pvalb, Tac2, Calb1, Nts,* and *Gpr83* across subdivisions of the MB. These markers occupied characteristic but partially overlapping spatial domains that broadly correspond to classical anatomical subdivisions, while also revealing molecular organization that extends across atlas-defined boundaries. We next identified distinct patterns of voltage-gated ion channel expression across MB neuronal populations, with transcripts encoding Na_V_1.1, K_V_7.2, K_V_7.3, and HCN1 showing differential enrichment across molecular populations and MB subregions. Finally, we demonstrated distinct intrinsic electrophysiological properties across MB subregions, with LM neurons exhibiting substantially lower excitability and different firing patterns compared with MnM neurons. Together, these findings establish a molecular framework linking neuronal population identity, spatial organization, and ion channel expression within the MB, and identify candidate channels that may contribute to differences in intrinsic excitability across MB neuronal populations. Given that several of these channels are encoded by genes implicated in NDDs, this framework also provides a foundation for future work investigating how altered ion channel function may affect MB circuits in neurological disease.

### Molecular organization of MB neuronal populations

Our previous scRNAseq analysis identified multiple transcriptionally distinct neuronal populations within the MB that are organized into characteristic spatial domains broadly corresponding to classical anatomical subdivisions (Mickelsen et al., 2020. Importantly, these transcriptomic populations were defined by combinatorial patterns of gene expression rather than by any single marker gene, and individual marker transcripts were enriched, but rarely restricted, to a given population. We therefore used FISH to determine how representative cluster-enriched markers are distributed across the rostrocaudal extent of the MB and how these molecular domains relate to atlas-defined anatomical boundaries. Overall, the spatial distributions of *Pvalb*, *Tac2*, *Calb1*, *Gpr83*, and *Nts* broadly aligned with their predicted anatomical enrichment, while also revealing overlapping expression patterns between neighboring subregions that are not obvious from the scRNAseq data alone. One example was *Nts* expression. Although our scRNAseq data identified *Nts* as most strongly enriched within the MM, FISH revealed an additional population of *Nts*-expressing neurons extending into the ML subdivision, particularly in rostral and middle portions of the MB before becoming increasingly restricted to the MM in more caudal sections. Similarly, *Pvalb* expression extended beyond the classically defined borders of the MnM (Paxinos, 2012), consistent with recent protein localization studies (Liu et al., 2025). Rather than indicating disagreement between transcriptomic and anatomical datasets, these observations likely reflect complex and graded patterns of transcript enrichment across neighboring neuronal populations and illustrate how spatial mapping complements transcriptomic clustering.

Overall, our findings are highly consistent with previous anatomical and molecular studies of MB organization. Classically, the MB are divided into the lateral and medial mammillary nuclei, with the latter further subdivided into the MnM, ML, and MM subdivisions (Seki and Zyo, 1984; Allen and Hopkins, 1988, 1989; Paxinos, 2012; Bubb et al., 2017; Żakowski and Zawistowski, 2023). Consistent with earlier immunohistochemical studies, we observed widespread *Calb1* expression across ventral regions of the medial nucleus and enrichment of *Pvalb* within more dorsal domains (Celio, 1990; Séquier et al., 1990; Rogers and Résibois, 1992; Liu et al., 2025, Li et al. 2025), further supporting molecular heterogeneity among neuronal populations within the MB. More broadly, the molecular populations and spatial organization identified here are generally consistent with recent studies employing single-cell transcriptomics (Huang et al, 2020; Li et al., 2025) and genetically defined subpopulations (Liu et al., 2025), which similarly reveal substantial molecular diversity within and across classical anatomical subdivisions.

The distribution of *Nts* neurons remains less well characterized. While both our scRNAseq and FISH data identify *Nts*-expressing neurons within the medial nucleus, previous immunohistochemical studies localized neurotensin primarily to fibers traversing the MB, suggesting that these fibers originate from subicular afferents rather than local MB neurons (Kahn et al., 1982; Kiyama et al., 1986). One possible explanation for this discrepancy is developmental regulation of *Nts* expression, as neurotensin immunoreactivity is prominent during early postnatal development but declines substantially in adulthood in both rodents and humans (Kiyama et al., 1986; Sakamoto et al., 1986, 1987; Langevin and Emson, 1982; Roberts et al., 1983; Mai et al., 1987). As our analyses were performed in juvenile mice (P30–35), developmental differences may partially account for these observations, although this possibility will require direct investigation.

Together, these findings demonstrate that transcriptomically defined MB neuronal populations are organized as overlapping molecular domains or gradients that broadly align with classical anatomical subdivisions while revealing previously unappreciated spatial heterogeneity in the distribution of cluster-enriched markers. This integrated molecular and anatomical framework reveals spatial heterogeneity that is not fully captured by classical anatomical subdivisions and provides a foundation for investigating how molecularly defined MB neuronal populations contribute to the functional organization of Papez circuit networks.

### Molecular organization of MB subcircuits and functional implications

The MB occupy a key position within the Papez circuit, receiving topographically organized input from the subiculum via the fornix and projecting to both the anterior thalamic nuclei (ATN) and the ventral tegmental nuclei of Gudden through the mammillothalamic (mtt) and mammillotegmental (mtg) tracts, respectively (Guillery, 1955; Watanabe and Kawana, 1980; Seki and Zyo, 1984; Hayakawa and Zyo, 1989, 1991; Allen and Hopkins, 1989; Shibata, 1992; Vann et al., 2007; Witter, 2006; Bienkowski et al., 2018). These projections are highly organized, with neurons of the ML projecting to the ventral anterior thalamic nucleus (AV), MM neurons projecting to the anteromedial nucleus (AM), and LM neurons projecting to the anterodorsal nucleus (AD) (Watanabe and Kawana, 1980; Seki and Zyo, 1984; Shibata, 1992; Mickelsen et al., 2020). Recent work further demonstrated that subicular inputs are similarly organized, with the dorsal MnM receiving the highest density of hippocampal input and exhibiting the strongest entrainment to hippocampal theta oscillations (Nitzan and Buzsáki, 2024).

Our molecular characterization provides a framework for relating these well-established anatomical pathways to genetically defined neuronal populations. In particular, the enrichment of *Pvalb* neurons within the dorsal MnM, consistent with recent work (Liu et al., 2025), suggests that this population may correspond to neurons receiving dense subicular input and participating in hippocampal theta-related activity (Nitzan and Buzsáki, 2024). Likewise, the distinct molecular composition of the LM, including *Pvalb*- and *Tac2*-expressing neurons, places these populations within the circuitry projecting to the AD (Watanabe and Kawana, 1980; Seki and Zyo, 1984; Shibata, 1992). Together, these findings provide molecular resolution to previously described MB subcircuits and establish genetically defined neuronal populations that can now be interrogated using modern circuit-based approaches.

Functionally, a significant body of evidence indicates that these anatomically distinct subcircuits make different contributions to spatial memory. The LM contains head direction neurons that project to the AD, another major component of the head direction network, and disruption of the LM impairs both head direction signaling and spatial working memory (Taube, 1995; Blair et al., 1998, 1999; Radyushkin et al., 2005; Bassett et al., 2007). In contrast, neurons within the MM modulate their activity according to running speed and angular head velocity (Sharp and Turner-Williams, 2005; Dillingham et al., 2024), lesions of the MM impair spatial working memory (Santín et al., 1999; Vann and Aggleton, 2003), and neurons within this region are entrained to hippocampal theta oscillations (Kocsis and Vertes, 1994; Bland et al., 1995; Kirk et al., 1996; Żakowski et al., 2017; Nitzan and Buzsáki, 2024). Furthermore, disruption of the fornix, which provides the principal hippocampal input to the MM, produces deficits in episodic memory in humans (McMackin et al., 1995; Tsivilis et al., 2008). More recent work has defined the role of distinct MM subcircuits in driving distinct patterns of locomotor behavior (Liu et al, 2025).

Although our data do not directly establish the functional roles of individual MB neuronal populations, they identify candidate cell populations that are positioned within these previously defined circuits. For example, the localization of *Pvalb* neurons to both the dorsal MnM and LM raises the possibility that genetically related neuronal populations participate in distinct aspects of MB function depending on their circuit connectivity. More broadly, the molecular framework established here provides a foundation for future studies linking genetically defined MB neuronal populations with their intrinsic electrophysiological properties, long-range connectivity, and contributions to spatial memory and episodic memory.

### Electrophysiological findings implicating differential expression of ion channels among MB subregions

Recordings from guinea pig MB show clear differences in excitability between MM and LM neurons. LM neurons display repetitive firing when held at depolarized potentials that switches to an all-or-none burst when held at hyperpolarized potentials (Llinás and Alonso, 1992). On the other hand, MM neurons fire in a rhythmic bursting pattern that is simply delayed when cells are held at hyperpolarized potentials (Alonso and Llinás, 1992). A recent study focused on *Calb1*- and *Pvalb*-positive neurons in the MM and found that they represent two electrophysiologically distinct populations within the same nucleus (Liu et al., 2025). These differences in firing properties likely reflect, at least in part, differential ion channel expression across MB neuronal populations. Our electrophysiological findings confirm distinct intrinsic properties between MB subregions (Fig. 11), while our molecular data identify candidate ion channels that may contribute to these differences. Of note, we showed that several ion channel transcripts are uniquely enriched within the MB relative to other subregions of the VPH. This includes multiple potassium channels of the voltage-gated, inward-rectifying and two-pore variety as well as ryanodine receptors. Several of these channels are likely to contribute to the distinct firing properties of MB neuronal subpopulations. Importantly, many are encoded by genes implicated in NDDs (Thapar et al., 2017; Ismail and Shapiro, 2019). Although the cognitive deficits associated with these disorders have primarily been attributed to dysfunction of cortical and hippocampal circuits, the established role of the MB in memory suggests that alterations in MB neuronal excitability should also be considered in future studies.

### Differential patterns of expression for NDD associated ion channels

Our work shows that ion channels belonging to the Na_V_, K_V_, and HCN families are differentially expressed within the MB. The patterns revealed by scRNAseq data were generally consistent with anatomical evidence provided by FISH, although some variability was evident for *Hcn* transcripts. In the case of *Hcn2*, the signal detected likely represents both neuronal and non-neuronal populations in the MB, as the scRNAseq showed it in both groups. However, it is unclear what may account for the inconsistency seen in *Hcn3,* as this transcript was not heavily detected in the scRNAseq. Still, FISH reveals strong expression throughout central portions of the MB. Interestingly, this expression pattern contrasts with work showing that *Hcn3* is most highly expressed during embryonic stages, being replaced by *Hcn1* and *Hcn2* postnatally (Schlusche et al., 2021). This discrepancy warrants further investigation, potentially at the protein level; however, for the purposes of this study, we focused on the ion channels more strongly expressed postnatally that are also associated with NDDs. More specifically, among the previously mentioned families, transcripts encoding Na_V_1.1, K_V_7.2, K_V_7.3 and HCN1 show some level of enrichment within specific neuronal subpopulations. Not only might these channels play key roles in determining the intrinsic firing properties of these neurons, but each of the aforementioned channels is commonly associated with NDDs (Deciphering Developmental Disorders Study, 2017; Simkin et al., 2022).

### Na_V_1.1 expression in the MB and implications

Pathogenic variants in *SCN1A*, which encodes the voltage-gated sodium channel Na_V_1.1, are a major cause of Dravet syndrome, a developmental and epileptic encephalopathy characterized by severe seizures and cognitive impairment (Mantegazza et al., 2021). Although the consequences of *SCN1A* dysfunction have largely been investigated in cortical and hippocampal circuits, our data demonstrate robust *Scn1a* expression across several MB neuronal populations, with particularly strong enrichment in *Pvalb*+ neurons of the MnM. This expression pattern identifies MB *Pvalb*+ neurons as a previously underappreciated population that may be affected by *SCN1A* dysfunction and suggests that Na_V_1.1 may contribute to their intrinsic excitability. Consistent with this possibility, conditional deletion of *Scn1a* from *Pvalb*-expressing neurons produces spatial memory deficits in addition to seizures (Tatsukawa et al., 2018). Together, these observations raise the possibility that altered function of MB *Pvalb*+ neurons may contribute to the cognitive phenotypes associated with *SCN1A*-related disorders, although this remains to be tested directly. More broadly, our findings identify additional MB neuronal populations expressing ion channels encoded by NDD-associated genes and provide a framework for investigating their potential contribution to disease-associated circuit dysfunction.

### K_V_7 expression in the MB and implications

K_V_7.2 and K_V_7.3 exhibit distinct expression patterns within the MB. These channels contribute to the M-current, a slowly activating, non-inactivating potassium current that limits repetitive firing and neuronal hyperexcitability (Wang et al., 1998). The M-current is associated with heteromeric channels containing both K_V_7.2 and K_V_7.3 subunits, but there are populations of neurons that express only one channel or the other (King et al., 2014; Goff and Goldberg, 2019; Sun et al., 2019). Our data suggest that the MB may also have populations of neurons expressing only one or the other subunit, as *Kcnq2* transcripts appear at moderate levels across the MB while *Kcnq3* is sparse but concentrated in the LM. Pathogenic variants in *KCNQ2* and *KCNQ3* are associated with neurodevelopmental disorders ranging from intellectual disability to developmental and epileptic encephalopathy (Nappi et al., 2020). Functional studies further demonstrate that subunit composition can influence the consequences of disease-associated variants (Abidi et al., 2015; Kim et al., 2018; Tran et al., 2020; Vanoye et al., 2022; Ye et al., 2023; Zhang et al., 2024). If the transcript distributions identified here are reflected in functional channel composition, disruption of K_V_7 signaling could therefore have different effects across MB neuronal populations. In particular, the co-expression of *Kcnq2* and *Kcnq3* within subsets of LM neurons raises the possibility that these neurons differ in their sensitivity to perturbations of K_V_7.2 or K_V_7.3 compared with medial MB populations, where *Kcnq2* predominates. These findings identify K_V_7 channels as candidate determinants of population-specific MB excitability and provide a framework for investigating how their dysfunction affects MB physiology.

### Changes in MB neuron excitability in neurodegenerative disorders

Studies of Alzheimer’s disease (AD) have increasingly implicated the MB in disease-associated cognitive dysfunction. Early neuropathological studies demonstrated Aβ accumulation and degeneration within the MB of AD patients (Grossi et al., 1989; Baloyannis et al., 2016). More recently, studies using the 5XFAD mouse model showed that Aβ deposition occurs in the MB early in disease progression and is accompanied by increased neuronal excitability (Canter et al., 2019). A subsequent study reported that changes in excitability and neuronal abundance were preferentially associated with the LM, with comparatively little change detected in the MM (Huang et al., 2023). Interpretation of the accompanying transcriptomic findings, however, is complicated by differences in the molecular annotation of MB neuronal populations across studies. The population designated as LM by Huang et al. expressed *Tac2*, *Prlr, Esr1, Nr4a2*, and *Pdyn*, several of which are also characteristic of *Kiss1*+ KNDy neurons in the arcuate nucleus (Campbell et al., 2017; Mickelsen et al., 2020; Yu et al., 2022). This overlap in molecular signatures makes it difficult to unambiguously relate the reported transcriptomic changes to the LM populations defined in our dataset or to identify candidate mechanisms underlying the electrophysiological phenotype. More recent work using the same 5XFAD model has also demonstrated hyperexcitability of MM neurons, potentially involving alterations in persistent sodium and T-type calcium currents (Stincic et al., 2025). Together, these studies suggest that altered excitability may extend across multiple MB neuronal populations during AD progression. Our molecular characterization of ion channel expression across these populations provides a framework for identifying candidate conductances that may contribute to these physiological changes.

Although the molecular basis of altered excitability in the LM remains unknown, our data identify several candidate ion channels that warrant further investigation. We found that transcripts encoding multiple voltage-gated potassium channels, including *Kcna2, Kcnc2, Kcnc4* and *Kcnq3*, are enriched in the LM relative to other MB subregions (Fig. 3). These channels regulate distinct aspects of intrinsic excitability, including action potential repolarization, high-frequency firing, and suppression of repetitive firing through the M-current (Wang et al., 1994; Brown and Passmore, 2009; Kaczmarek and Zhang, 2017). Alterations in the expression or function of any of these channels could therefore contribute to the increased excitability reported in LM neurons. Similarly, our data show that medial MB subregions express multiple Na_V_ channels together with the T-type calcium channel *Cacna1g*, each of which could influence neuronal excitability if altered in AD. Because these channels are also expressed in the LM, they likewise represent candidates for mediating the physiological changes observed in this region. Together, our findings provide a molecular framework for investigating the ion channel mechanisms that underlie altered MB excitability in both neurodegenerative and neurodevelopmental disorders.

### Limitations and future directions

Although our study links the molecular identity of MB neuronal populations with the spatial distribution of voltage-gated ion channel transcripts, several important questions remain. First, because our analyses were performed primarily at the transcript level, future studies will be needed to determine whether these expression patterns are reflected in channel protein abundance and localization. Immunohistochemical approaches may help address this question, although the dense network of afferent and efferent fibers within the MB may complicate assignment of channel protein to specific neuronal populations and subcellular compartments. Second, while our electrophysiological recordings demonstrate distinct intrinsic properties of neurons across MB subregions, the specific conductances underlying these differences remain unknown. Pharmacological or genetic manipulation of candidate ion channels identified here will be necessary to establish their contributions to MB firing properties. Finally, although several of the channels examined are encoded by genes implicated in NDDs, our study does not directly address how their dysfunction affects MB neurons or contributes to disease phenotypes. Extending these analyses to well-characterized disease models carrying variants in genes such as *Scn1a* or *Kcnq2*, together with molecular, electrophysiological and imaging approaches, will help determine how channel dysfunction alters MB circuit function. Together, our findings establish a molecular and anatomical framework for understanding ion channel expression across MB neuronal populations and provide a foundation for determining how these differences shape the functional organization of MB circuits in health and disease.

## Data availability

The data that support the findings of this study are available from the corresponding authors upon reasonable request.

## Acknowledgments

We gratefully acknowledge all members of the Jackson and Tzingounis labs for support, assistance, and helpful discussions and C. O’Connell for imaging support. We also acknowledge R. Downing for assistance and discussion. Project supported by the National Institutes of Health grant R01MH112739 (to A.C.J.), as well as National Institute of Neurological Disorders and Stroke NS101596 and National Heart, Lung, and Blood Institute HL137094 grants (to A.V.T.), and an NIH Shared Instrumentation Grant S10OD016435 (to A. Nishiyama) for imaging support.

## REFERENCES

Abidi A, Devaux JJ, Molinari F, Alcaraz G, Michon FX, Sutera-Sardo J, Becq H, Lacoste C, Altuzarra C, Afenjar A, Mignot C, Doummar D, Isidor B, Guyen SN, Colin E, De La Vaissière S, Haye D, Trauffler A, Badens C, Prieur F, Lesca G, Villard L, Milh M, Aniksztejn L. A recurrent KCNQ2 pore mutation causing early onset epileptic encephalopathy has a moderate effect on M current but alters subcellular localization of Kv7 channels. Neurobiol Dis. 2015 Aug;80:80–92. doi: 10.1016/j.nbd.2015.04.017. Epub 2015 May 22. PMID: 26007637.

Aggleton JP, Brown MW. Episodic memory, amnesia, and the hippocampal-anterior thalamic axis. Behav Brain Sci. 1999 Jun;22(3):425–44; discussion 444-89. PMID: 11301518.

Aggleton JP, O’Mara SM, Vann SD, Wright NF, Tsanov M, Erichsen JT. Hippocampal-anterior thalamic pathways for memory: uncovering a network of direct and indirect actions. Eur J Neurosci. 2010 Jun;31(12):2292–307. doi: 10.1111/j.1460-9568.2010.07251.x. Epub 2010 Jun 14. PMID: 20550571; PMCID: PMC2936113.

Allen GV, Hopkins DA. Mamillary body in the rat: a cytoarchitectonic, Golgi, and ultrastructural study. J Comp Neurol. 1988 Sep 1;275(1):39–64. doi:10.1002/cne.902750105. PMID: 3139720.

Allen GV, Hopkins DA. Mamillary body in the rat: topography and synaptology of projections from the subicular complex, prefrontal cortex, and midbrain tegmentum. J Comp Neurol. 1989;286:311–336. doi: 10.1002/cne.902860303.

Alonso A, Llinás RR. Electrophysiology of the mammillary complex in vitro. II. Medial mammillary neurons. J Neurophysiol. 1992 Oct;68(4):1321–31. doi: 10.1152/jn.1992.68.4.1321. PMID: 1432086.

Arts NJM, Pitel AL, Kessels RPC. The contribution of mamillary body damage to Wernicke’s encephalopathy and Korsakoff’s syndrome. Handb Clin Neurol. 2021;180:455–475. doi: 10.1016/B978-0-12-820107-7.00029-X. PMID: 34225949.

Baloyannis SJ, Mavroudis I, Baloyannis IS, Costa VG. Mammillary Bodies in Alzheimer’s Disease: A Golgi and Electron Microscope Study. Am J Alzheimers Dis Other Demen. 2016 May;31(3):247–56. doi: 10.1177/1533317515602548. Epub 2015 Sep 22. PMID: 26399484.

Balschun D, Wolfer DP, Bertocchini F, Barone V, Conti A, Zuschratter W, Missiaen L, Lipp HP, Frey JU, Sorrentino V. Deletion of the ryanodine receptor type 3 (RyR3) impairs forms of synaptic plasticity and spatial learning. EMBO J. 1999 Oct 1;18(19):5264–73. doi: 10.1093/emboj/18.19.5264. PMID: 10508160; PMCID: PMC1171597.

Bassett JP, Tullman ML, Taube JS. Lesions of the tegmentomammillary circuit in the head direction system disrupt the head direction signal in the anterior thalamus. J Neurosci. 2007 Jul 11;27(28):7564–77. doi:10.1523/JNEUROSCI.0268-07.2007. PMID: 17626218; PMCID: PMC6672597.

Bean BP. The action potential in mammalian central neurons. Nat Rev Neurosci. 2007 Jun;8(6):451–65. doi: 10.1038/nrn2148. PMID: 17514198.

Biel M, Wahl-Schott C, Michalakis S, Zong X. Hyperpolarization-activated cation channels: from genes to function. Physiol Rev. 2009 Jul;89(3):847–85. doi: 10.1152/physrev.00029.2008. PMID: 19584315.

Bienkowski MS, Bowman I, Song MY, Gou L, Ard T, Cotter K, Zhu M, Benavidez NL, Yamashita S, Abu-Jaber J, Azam S, Lo D, Foster NN, Hintiryan H, Dong HW. Integration of gene expression and brain-wide connectivity reveals the multiscale organization of mouse hippocampal networks. Nat Neurosci. 2018;21:1628–1643. doi: 10.1038/s41593-018-0241-y.

Blair HT, Cho J, Sharp PE. Role of the lateral mammillary nucleus in the rat head direction circuit: a combined single unit recording and lesion study. Neuron. 1998 Dec;21(6):1387–97. doi: 10.1016/s0896-6273(00)80657-1. Erratum in: Neuron 1999 Jan;22(1):199. PMID: 9883731.

Blair HT, Cho J, Sharp PE. The anterior thalamic head-direction signal is abolished by bilateral but not unilateral lesions of the lateral mammillary nucleus. J Neurosci. 1999 Aug 1;19(15):6673–83. doi:0.1523/JNEUROSCI.19-15-06673.1999. PMID: 10414996; PMCID: PMC6782818.

Bland BH, Konopacki J, Kirk IJ, Oddie SD, Dickson CT. Discharge patterns of hippocampal theta-related cells in the caudal diencephalon of the urethan-anesthetized rat. J Neurophysiol. 1995 Jul;74(1):322–33. doi:10.1152/jn.1995.74.1.322. PMID: 7472334.

Branco T, Tozer A, Magnus CJ, Sugino K, Tanaka S, Lee AK, Wood JN, Sternson SM. Near-Perfect Synaptic Integration by Na_V_1.7 in Hypothalamic Neurons Regulates Body Weight. Cell. 2016 Jun 16;165(7):1749–1761. doi: 10.1016/j.cell.2016.05.019. PMID: 27315482; PMCID: PMC4912688.

Brickley SG, Aller MI, Sandu C, Veale EL, Alder FG, Sambi H, Mathie A, Wisden W. TASK-3 two-pore domain potassium channels enable sustained high-frequency firing in cerebellar granule neurons. J Neurosci. 2007 Aug 29;27(35):9329–40. doi: 10.1523/JNEUROSCI.1427-07.2007. PMID: 17728447; PMCID: PMC6673138.

Brown DA, Passmore GM. Neural KCNQ (Kv7) channels. Br J Pharmacol. 2009 Apr;156(8):1185–95. doi:10.1111/j.1476-5381.2009.00111.x. Epub 2009 Mar 9. PMID: 19298256; PMCID: PMC2697739.

Bubb EJ, Kinnavane L, Aggleton JP. Hippocampal -diencephalic -cingulate networks for memory and emotion: An anatomical guide. Brain Neurosci Adv. 2017 Aug 4;1(1):2398212817723443. doi: 10.1177/2398212817723443. PMID: 28944298; PMCID: PMC5608081.

Callen DJ, Black SE, Gao F, Caldwell CB, Szalai JP. Beyond the hippocampus: MRI volumetry confirms widespread limbic atrophy in AD. Neurology. 2001 Nov 13;57(9):1669–74. doi: 10.1212/wnl.57.9.1669. PMID: 11706109.

Campbell JN, Macosko EZ, Fenselau H, Pers TH, Lyubetskaya A, Tenen D, Goldman M, Verstegen AM, Resch JM, McCarroll SA, Rosen ED, Lowell BB, Tsai LT. A molecular census of arcuate hypothalamus and median eminence cell types. Nat Neurosci. 2017 Mar;20(3):484–496. doi: 10.1038/nn.4495. Epub 2017 Feb 6. PMID: 28166221; PMCID: PMC5323293.

Catterall WA. Voltage gated sodium and calcium channels: Discovery, structure, function, and Pharmacology. Channels (Austin). 2023 Dec;17(1):2281714. doi: 10.1080/19336950.2023.2281714. Epub 2023 Nov 20. PMID: 37983307; PMCID: PMC10761118.

Celio MR. Calbindin D-28k and parvalbumin in the rat nervous system. Neuroscience. 1990;35:375–475. doi: 10.1016/0306-4522(90)90091-h.

Copenhaver, B. R. et al. The fornix and mammillary bodies in older adults with Alzheimer’s disease, mild cognitive impairment, and cognitive complaints: A volumetric MRI study. Psychiatry Res. -Neuroimaging 147, 93–103 (2006).

Deciphering Developmental Disorders Study. Prevalence and architecture of de novo mutations in developmental disorders. Nature. 2017 Feb 23;542(7642):433–438. doi: 10.1038/nature21062. Epub 2017Jan 25. PMID: 28135719; PMCID: PMC6016744.

Dillingham CM, Frizzati A, Nelson AJ, Vann SD. How do mammillary body inputs contribute to anterior thalamic function? Neurosci Biobehav Rev. 2015 Jul;54:108–19. doi: 10.1016/j.neubiorev.2014.07.025. Epub 2014 Aug 11. PMID: 25107491; PMCID: PMC4462591

Dillingham CM, Wilson JJ, Vann SD. Electrophysiological Properties of the Medial Mammillary Bodies across the Sleep-Wake Cycle. eNeuro. 2024 Apr 26;11(4):ENEURO.0447-23.2024. doi: 10.1523/ENEURO.0447-23.2024. PMID: 38621991; PMCID: PMC11055652.

Enyedi P, Czirják G. Molecular background of leak K+ currents: two-pore domain potassium channels. Physiol Rev. 2010 Apr;90(2):559–605. doi: 10.1152/physrev.00029.2009. PMID: 20393194.

Gail Canter R, Huang WC, Choi H, Wang J, Ashley Watson L, Yao CG, Abdurrob F, Bousleiman SM, Young JZ, Bennett DA, Delalle I, Chung K, Tsai LH. 3D mapping reveals network-specific amyloid progression and subcortical susceptibility in mice. Commun Biol. 2019 Oct 4;2:360. doi: 10.1038/s42003-019-0599-8. PMID:31602409; PMCID: PMC6778135.

Galeotti N, Quattrone A, Vivoli E, Norcini M, Bartolini A, Ghelardini C. Different involvement of type 1, 2, and 3 ryanodine receptors in memory processes. Learn Mem. 2008 Apr 25;15(5):315–23. doi: 10.1101/lm.929008. PMID: 18441289; PMCID: PMC2364603.

Goff KM, Goldberg EM. Vasoactive intestinal peptide-expressing interneurons are impaired in a mouse model of Dravet syndrome. Elife. 2019 Jul 8;8:e46846. doi: 10.7554/eLife.46846. PMID: 31282864; PMCID: PMC6629374.

Grossi D, Lopez OL, Martinez AJ. Mamillary bodies in Alzheimer’s disease. Acta Neurol Scand. 1989 Jul;80(1):41–5. doi: 10.1111/j.1600-0404.1989.tb03840.x. PMID: 2506728.

Guillery RW. A quantitative study of the mamillary bodies and their connexions. J Anat. 1955;89:19–32.

Hallmann K, Durner M, Sander T, Steinlein OK. Mutation analysis of the inwardly rectifying K(+) channels KCNJ6 (GIRK2) and KCNJ3 (GIRK1) in juvenile myoclonic epilepsy. Am J Med Genet. 2000 Feb 7;96(1):8–11. doi: 10.1002/(sici)1096-8628(20000207)96:1<8::aid-ajmg3>3.0.co;2-s. PMID: 10686544.

Hayakawa T, Zyo K. Quantitative and ultrastructural study of ascending projections to the medial mammillary nucleus in the rat. Anat Embriol. 1991;184:611–622. doi: 10.1007/BF00942583.

Hayakawa T, Zyo K. Retrograde double-labelling study of the mammillothalamic and the mammillotegmental projections in the rat. J Comp Neurol. 1989;284:1–11. doi: 10.1002/cne.902840102.

Hibino H, Inanobe A, Furutani K, Murakami S, Findlay I, Kurachi Y. Inwardly rectifying potassium channels: their structure, function, and physiological roles. Physiol Rev. 2010 Jan;90(1):291–366. doi: 10.1152/physrev.00021.2009. PMID: 20086079.

Huang WC, Peng Z, Murdock MH, Liu L, Mathys H, Davila-Velderrain J, Jiang X, Chen M, Ng AP, Kim T, Abdurrob F, Gao F, Bennett DA, Kellis M, Tsai LH. Lateral mammillary body neurons in mouse brain are disproportionately vulnerable in Alzheimer’s disease. Sci Transl Med. 2023 Apr 19;15(692):eabq1019. doi: 10.1126/scitranslmed.abq1019. Epub 2023 Apr 19. PMID: 37075128; PMCID: PMC10511020.

Ismail FY, Shapiro BK. What are neurodevelopmental disorders? Curr Opin Neurol. 2019 Aug;32(4):611–616. doi: 10.1097/WCO.0000000000000710. PMID: 31116115.

Jan LY, Jan YN. Voltage-gated potassium channels and the diversity of electrical signalling. J Physiol. 2012 Jun 1;590(11):2591–9. doi: 10.1113/jphysiol.2011.224212. Epub 2012 Mar 19. PMID: 22431339; PMCID: PMC3424718.

Jankowski MM, Ronnqvist KC, Tsanov M, Vann SD, Wright NF, Erichsen JT, Aggleton JP, O’Mara SM. The anterior thalamus provides a subcortical circuit supporting memory and spatial navigation. Front Syst Neurosci. 2013 Aug 30;7:45. doi: 10.3389/fnsys.2013.00045. PMID: 24009563; PMCID: PMC3757326.

Kaczmarek LK, Zhang Y. Kv3 Channels: Enablers of Rapid Firing, Neurotransmitter Release, and Neuronal Endurance. Physiol Rev. 2017 Oct 1;97(4):1431–1468. doi: 10.1152/physrev.00002.2017. PMID: 28904001;PMCID: PMC6151494.

Kahn D, Hou-Yu A, Zimmerman EA. Localization of neurotensin in the hypothalamus. Ann NY Acad Sci. 1982;400:117–131. doi: 10.1111/j.1749-6632.1982.tb31564.x.

Kessi M, Peng J, Duan H, He H, Chen B, Xiong J, Wang Y, Yang L, Wang G, Kiprotich K, Bamgbade OA, He F, Yin F. The Contribution of HCN Channelopathies in Different Epileptic Syndromes, Mechanisms, Modulators, and Potential Treatment Targets: A Systematic Review. Front Mol Neurosci. 2022 May 19;15:807202. doi: 10.3389/fnmol.2022.807202. PMID: 35663267; PMCID: PMC9161305.

Kim EC, Zhang J, Pang W, Wang S, Lee KY, Cavaretta JP, Walters J, Procko E, Tsai NP, Chung HJ. Reduced axonal surface expression and phosphoinositide sensitivity in K_v_7 channels disrupts their function to inhibit neuronal excitability in Kcnq2 epileptic encephalopathy. Neurobiol Dis. 2018 Oct;118:76–93. doi: 10.1016/j.nbd.2018.07.004. Epub 2018 Jul 6. PMID: 30008368; PMCID: PMC6415549.

King CH, Lancaster E, Salomon D, Peles E, Scherer SS. Kv7.2 regulates the function of peripheral sensory neurons. J Comp Neurol. 2014 Oct 1;522(14):3262–80. doi: 10.1002/cne.23595. Epub 2014 Apr 12. PMID: 24687876; PMCID: PMC4428907.

Kirk IJ, Oddie SD, Konopacki J, Bland BH. Evidence for differential control of posterior hypothalamic, supramammillary, and medial mammillary theta-related cellular discharge by ascending and descending pathways. J Neurosci. 1996 Sep 1;16(17):5547–54. doi: 10.1523/JNEUROSCI.16-17-05547.1996. PMID:8757266; PMCID: PMC6578895.

Kiyama H, Shiosaka S, Sakamoto N, Michel JP, Pearson J, Tohyama M. A neurotensin-immunoreactive pathway from the subiculum to the mammillary body in the rat. Brain Res. 1986;375:357–359. doi: 10.1016/0006-8993(86)90757-2.

Kocsis B, Vertes RP. Characterization of neurons of the supramammillary nucleus and mammillary body that discharge rhythmically with the hippocampal theta rhythm in the rat. J Neurosci. 1994 Nov;14(11 Pt 2):7040–52. doi: 10.1523/JNEUROSCI.14-11-07040.1994. PMID: 7965097; PMCID: PMC6577300.

Kril JJ, Harper CG. Neuroanatomy and neuropathology associated with Korsakoff’s syndrome. Neuropsychol Rev. 2012 Jun;22(2):72–80. doi: 10.1007/s11065-012-9195-0. Epub 2012 Apr 14. PMID: 22528862; PMCID: PMC3371089.

Langevin H, Emson PC. Distribution of substance P, somatostatin and neurotensin in the human hypothalamus. Brain Res. 1982;246:65–69. doi: 10.1016/0006-8993(82)90142-1.

Lein ES, Hawrylycz MJ, Ao N, Ayres M, Bensinger A, Bernard A, Boe AF, Boguski MS, Brockway KS, Byrnes EJ, Chen L, Chen L, Chen TM, Chin MC, Chong J, Crook BE, Czaplinska A, Dang CN, Datta S, Dee NR, Desaki AL, Desta T, Diep E, Dolbeare TA, Donelan MJ, Dong HW, Dougherty JG, Duncan BJ, Ebbert AJ, Eichele G, Estin LK, Faber C, Facer BA, Fields R, Fischer SR, Fliss TP, Frensley C, Gates SN, Glattfelder KJ, Halverson KR, Hart MR, Hohmann JG, Howell MP, Jeung DP, Johnson RA, Karr PT, Kawal R, Kidney JM, Knapik RH, Kuan CL, Lake JH, Laramee AR, Larsen KD, Lau C, Lemon TA, Liang AJ, Liu Y, Luong LT, Michaels J, Morgan JJ, Morgan RJ, Mortrud MT, Mosqueda NF, Ng LL, Ng R, Orta GJ, Overly CC, Pak TH, Parry SE, Pathak SD, Pearson OC, Puchalski RB, Riley ZL, Rockett HR, Rowland SA, Royall JJ, Ruiz MJ, Sarno NR, Schaffnit K, Shapovalova NV, Sivisay T, Slaughterbeck CR, Smith SC, Smith KA, Smith BI, Sodt AJ, Stewart NN, Stumpf KR, Sunkin SM, Sutram M, Tam A, Teemer CD, Thaller C, Thompson CL, Varnam LR, Visel A, Whitlock RM, Wohnoutka PE, Wolkey CK, Wong VY, Wood M, Yaylaoglu MB, Young RC, Youngstrom BL, Yuan XF, Zhang B, Zwingman TA, Jones AR. Genome-wide atlas of gene expression in the adult mouse brain. Nature. 2007 Jan 11;445(7124):168–76. doi: 10.1038/nature05453. Epub 2006 Dec 6. PMID: 17151600.

Li L, Guo Y, Jing W, Tang X, Zeng J, Hou Z, Song Y, He A, Li H, Zhu LQ, Lu Y, Li X. (2025). Cell-Type Specific Circuits in the Mammillary Body for Place and Object Recognition Memory. Adv Sci (Weinh). 12(13):e2409397. PMID: 39928529

Liu H, Shi Y, Zhang Q, Yue M, Qi Y, Xu B, Jing J, Zhang L, Yang K, Zheng M, Zhou J, Lu J, Gong L, He M. Two distinct cell types of the medial mammillary body forming segregated subcircuits. Mol Psychiatry. 2025 Jun 23. doi: 10.1038/s41380-025-03079-w. Epub ahead of print. PMID: 40550863.

Llinás RR, Alonso A. Electrophysiology of the mammillary complex in vitro. I. Tuberomammillary and lateral mammillary neurons. J Neurophysiol. 1992 Oct;68(4):1307–20. doi: 10.1152/jn.1992.68.4.1307. PMID: 1279134.

Mai JK, Triepel J, Metz J. Neurotensin in the human brain. Neuroscience. 1987;22:499–524. doi: 10.1016/0306-4522(87)90349-6.

Mantegazza M, Cestèle S, Catterall WA. Sodium channelopathies of skeletal muscle and brain. Physiol Rev. 2021 Oct 1;101(4):1633–1689. doi: 10.1152/physrev.00025.2020. Epub 2021 Mar 26. PMID: 33769100; PMCID: PMC8989381.

Marini C, Porro A, Rastetter A, Dalle C, Rivolta I, Bauer D, Oegema R, Nava C, Parrini E, Mei D, Mercer C, Dhamija R, Chambers C, Coubes C, Thévenon J, Kuentz P, Julia S, Pasquier L, Dubourg C, Carré W, Rosati A, Melani F, Pisano T, Giardino M, Innes AM, Alembik Y, Scheidecker S, Santos M, Figueiroa S, Garrido C, Fusco C, Frattini D, Spagnoli C, Binda A, Granata T, Ragona F, Freri E, Franceschetti S, Canafoglia L, Castellotti B, Gellera C, Milanesi R, Mancardi MM, Clark DR, Kok F, Helbig KL, Ichikawa S, Sadler L, Neupauerová J, Laššuthova P, Šterbová K, Laridon A, Brilstra E, Koeleman B, Lemke JR, Zara F, Striano P, Soblet J, Smits G, Deconinck N, Barbuti A, DiFrancesco D, LeGuern E, Guerrini R, Santoro B, Hamacher K, Thiel G, Moroni A, DiFrancesco JC, Depienne C. HCN1 mutation spectrum: from neonatal epileptic encephalopathy to benign generalized epilepsy and beyond. Brain. 2018 Nov 1;141(11):3160–3178. doi: 10.1093/brain/awy263. PMID: 30351409.

Matsuo N, Tanda K, Nakanishi K, Yamasaki N, Toyama K, Takao K, Takeshima H, Miyakawa T. Comprehensive behavioral phenotyping of ryanodine receptor type 3 (RyR3) knockout mice: decreased social contact duration in two social interaction tests. Front Behav Neurosci. 2009 May 7;3:3. doi: 10.3389/neuro.08.003.2009. PMID: 19503748; PMCID: PMC2691151.

McMackin D, Cockburn J, Anslow P, Gaffan D. Correlation of fornix damage with memory impairment in six cases of colloid cyst removal. Acta Neurochir. 1995;135:12–18. doi: 10.1007/BF02307408.

Mickelsen LE, Flynn WF, Springer K, Wilson L, Beltrami EJ, Bolisetty M, Robson P, Jackson AC. Cellular taxonomy and spatial organization of the murine ventral posterior hypothalamus. Elife. 2020 Oct 29;9:e58901. doi: 10.7554/eLife.58901. PMID: 33119507; PMCID: PMC7595735.

Nappi P, Miceli F, Soldovieri MV, Ambrosino P, Barrese V, Taglialatela M. Epileptic channelopathies caused by neuronal Kv7 (KCNQ) channel dysfunction. Pflugers Arch. 2020 Jul;472(7):881–898. doi: 10.1007/s00424-020-02404-2. Epub 2020 Jun 6. PMID: 32506321.

Nitzan N, Buzsáki G. Physiological characteristics of neurons in the mammillary bodies align with topographical organization of subicular inputs. Cell Rep. 2024 Aug 27;43(8):114539. doi: 10.1016/j.celrep.2024.114539. Epub 2024 Jul 23. PMID: 39052483; PMCID: PMC11475818.

Paxinos G. 2012. Paxinos and Franklin’s the Mouse Brain in Stereotaxic Coordinates. Amsterdam: Academic Press.

Radyushkin K, Anokhin K, Meyer BI, Jiang Q, Alvarez-Bolado G, Gruss P. Genetic ablation of the mammillary bodies in the Foxb1 mutant mouse leads to selective deficit of spatial working memory. Eur J Neurosci. 2005 Jan;21(1):219–29. doi:10.1111/j.1460-9568.2004.03844.x. PMID: 15654859.

Roberts GW, Allen Y, Crow TJ, Polak JM. Immunocytochemical localization on neuropeptides in the fornix of rat, monkey and man. Brain Res. 1983;263:151–155. doi: 10.1016/0006-8993(83)91213-1.

Rogers JH, Résibois A. Calretinin and calbindin-D28k in rat brain: patterns of partial co-localization. Neuroscience. 1992;51:843–865. doi: 10.1016/0306-4522(92)90525-7.

Sakamoto N, Michel JP, Kiyama H, Tohyama M, Kopp N, Pearson J. Neurotensin immunoreactivity in the human cingulate gyrus, hippocampal subiculum and mammillary bodies. Its potential role in memory processing. Brain Res. 1986;375:351–356. doi: 10.1016/0006-8993(86)90756-0.

Sakamoto N, Michel JP, Kopp N, Pearson J. Neurotensin immunoreactive neurons in the human infant diencephalon. Brain Res. 1987 Feb 10;403(1):31–42. doi: 10.1016/0006-8993(87)90119-3. PMID: 3548888.

Santín LJ, Rubio S, Begega A, Arias JL. Effects of mammillary body lesions on spatial reference and working memory tasks. Behav Brain Res. 1999 Jul;102(1-2):137–50. doi: 10.1016/s0166-4328(99)00011-x. PMID: 10403022.

Saper CB, German DC. Hypothalamic pathology in Alzheimer’s disease. Neurosci Lett. 1987 Mar 9;74(3):364–70. doi: 10.1016/0304-3940(87)90325-9. PMID: 2436113.

Schindelin J, Arganda-Carreras I, Frise E, Kaynig V, Longair M, Pietzsch T, Preibisch S, Rueden C, Saalfeld S, Schmid B, Tinevez JY, White DJ, Hartenstein V, Eliceiri K, Tomancak P, Cardona A. Fiji: an open-source platform for biological-image analysis. Nat Methods. 2012 Jun 28;9(7):676–82. doi: 10.1038/nmeth.2019. PMID: 22743772.

Schlusche AK, Vay SU, Kleinenkuhnen N, Sandke S, Campos-Martín R, Florio M, Huttner W, Tresch A, Roeper J, Rueger MA, Jakovcevski I, Stockebrand M, Isbrandt D. Developmental HCN channelopathy results in decreased neural progenitor proliferation and microcephaly in mice. Proc Natl Acad Sci U S A. 2021 Aug 31;118(35):e2009393118. doi: 10.1073/pnas.2009393118. PMID: 34429357; PMCID: PMC8536352.

Schwarz JR, Glassmeier G, Cooper EC, Kao TC, Nodera H, Tabuena D, Kaji R, Bostock H. KCNQ channels mediate IKs, a slow K+ current regulating excitability in the rat node of Ranvier. J Physiol. 2006 May 15;573(Pt 1):17–34. doi: 10.1113/jphysiol.2006.106815. Epub 2006 Mar 9. PMID: 16527853; PMCID: PMC1779690

Seki M, Zyo K. Anterior thalamic afferents from the mamillary body and the limbic cortex in the rat. J Comp Neurol. 1984;229:242–256. doi: 10.1002/cne.902290209.

Séquier JM, Hunziker W, Andressen C, Celio MR. Calbindin D-28k protein and mRNA localization in the rat brain. Eur J Neurosci. 1990;2:1118–1126. doi: 10.1111/j.1460-9568.1990.tb00023.x

Sharp PE, Turner-Williams S. Movement-related correlates of single-cell activity in the medial mammillary nucleus of the rat during a pellet-chasing task. J Neurophysiol. 2005 Sep;94(3):1920–7. doi: 10.1152/jn.00194.2005. Epub 2005 Apr 27. Erratum in: J Neurophysiol. 2005 Dec;94(6):4554. Turner-Williams, S [added]. PMID: 15857969.

Shibata H. Topographic organization of subcortical projections to the anterior thalamic nuclei in the rat. J Comp Neurol. 1992;323:117–127. doi: 10.1002/cne.903230110.

Shin W, Kweon H, Kang R, Kim D, Kim K, Kang M, Kim SY, Hwang SN, Kim JY, Yang E, Jim H, Kim E. *Scn2a* Haploinsufficiency in Mince Suppresses Hippocampal Neuronal Excitability, Excitatory Synaptic Drive and Long-Term Potentiation, and Spatial Learning and Memory. Front Mol Neurosci. 2019 June 4;12:145. doi:10.3389/fnmol.2019. PMID: 31249508

Simkin D, Ambrosi C, Marshall KA, Williams LA, Eisenberg J, Gharib M, Dempsey GT, George AL Jr, McManus OB, Kiskinis E. ’Channeling’ therapeutic discovery for epileptic encephalopathy through iPSC technologies. Trends Pharmacol Sci. 2022 May;43(5):392–405. doi: 10.1016/j.tips.2022.03.001. PMID: 35427475; PMCID: PMC9119009.

Singh NA, Charlier C, Stauffer D, DuPont BR, Leach RJ, Melis R, Ronen GM, Bjerre I, Quattlebaum T, Murphy JV, McHarg ML, Gagnon D, Rosales TO, Peiffer A, Anderson VE, Leppert M. A novel potassium channel gene, KCNQ2, is mutated in an inherited epilepsy of newborns. Nat Genet. 1998 Jan;18(1):25–9. doi: 10.1038/ng0198-25. PMID: 9425895.

Stincic TL, Qiu J, Martinson C, Kelly MJ, Ronnekleiv OK. Early onset of hyperexcitability of medial mammillary body neurons in 5XFAD^+^ mice-an early target of Alzheimer’s disease. J Neurophysiol. 2025 Sep 1;134(3):928–939. doi: 10.1152/jn.00310.2025. Epub 2025 Aug 6. PMID: 40767842; PMCID: PMC12453389.

Sun H, Lin AH, Ru F, Patil MJ, Meeker S, Lee LY, Undem BJ. KCNQ/M-channels regulate mouse vagal bronchopulmonary C-fiber excitability and cough sensitivity. JCI Insight. 2019 Mar 7;4(5):e124467. doi: 10.1172/jci.insight.124467. PMID: 30721152; PMCID: PMC6483509.

Tatsukawa T, Ogiwara I, Mazaki E, Shimohata A, Yamakawa K. Impairments in social novelty recognition and spatial memory in mice with conditional deletion of Scn1a in parvalbumin-expressing cells. Neurobiol Dis. 2018 Apr;112:24–34. doi:10.1016/j.nbd.2018.01.009. Epub 2018 Jan 11. PMID: 29337050.

Taube JS. Head direction cell, recorded in the anterior thalamic nuclei of freely moving rats. J. Neuorosci. 1995;15:70–86. doi: 10.1523/JNEUROSCI.15-01-00070.1995.

Thapar A, Cooper M, Rutter M. Neurodevelopmental disorders. Lancet Psychiatry. 2017 Apr;4(4):339-346. doi: 10.1016/S2215-0366(16)30376-5. Epub 2016 Dec 13. PMID: 27979720.

Tran B, Ji ZG, Xu M, Tsuchida TN, Cooper EC. Two *KCNQ2* Encephalopathy Variants in the Calmodulin-Binding Helix A Exhibit Dominant-Negative Effects and Altered PIP_2_ Interaction. Front Physiol. 2020 Sep 11;11:1144. doi: 10.3389/fphys.2020.571813. PMID: 33041849; PMCID: PMC7518097.

Tsivilis D, Vann SD, Denby C, Roberts N, Mayes AR, Montaldi D, Aggleton JP. A disproportionate role for the fornix and mammillary bodies in recall versus recognition memory. Nat Neurosci. 2008 Jul;11(7):834–42. doi: 10.1038/nn.2149. Epub 2008 Jun 15. PMID: 18552840.

Tsivilis D, Vann SD, Denby C, Roberts N, Mayes AR, Montaldi D, Aggleton JP. The importance of the fornix and mammillary bodies for human memory: a disproportionate role for recall versus recognition. Nat. Neurosci. 2008;11:834–842. doi: 10.1038/nn.2149.

Vacher H, Mohapatra DP, Trimmer JS. Localization and targeting of voltage-dependent ion channels in mammalian central neurons. Physiol Rev. 2008 Oct;88(4):1407–47. doi: 10.1152/physrev.00002.2008. PMID: 18923186; PMCID: PMC2587220.

Vann SD, Aggleton JP. Evidence of a spatial encoding deficit in rats with lesions of the mammillary bodies or mammillothalamic tract. J Neurosci. 2003 Apr 15;23(8):3506–14. doi: 10.1523/JNEUROSCI.23-08-03506.2003. PMID: 12716960; PMCID: PMC6742300.

Vann SD, Aggleton JP. The mammillary bodies: two memory systems in one? Nat Rev Neurosci. 2004 Jan;5(1):35–44. doi: 10.1038/nrn1299. PMID: 14708002.

Vann SD, Nelson AJ. The mammillary bodies and memory: more than a hippocampal relay. Prog Brain Res. 2015;219:163–85. doi: 10.1016/bs.pbr.2015.03.006. Epub 2015 May 16. PMID: 26072239; PMCID: PMC4498492.

Vann SD, Saunders RC, Aggleton JP. Distinct, parallel pathways link the medial mammillary bodies to the anterior thalamus in macaque monkeys. Eur J Neurosci. 2007;26:1575–1586. doi: 10.1111/j.1460-9568.2007.05773.x.

Vann SD. Re-evaluating the role of the mammillary bodies in memory. Neuropsychologia. 2010 Jul;48(8):2316–27. doi: 10.1016/j.neuropsychologia.2009.10.019. Epub 2009 Oct 30. PMID: 19879886.

Vann SD. Dismantling the Papez circuit for memory in rats. Elife. 2013 Jun 25;2:e00736. doi: 10.7554/eLife.00736. PMID: 23805381; PMCID: PMC3691571.

Vanoye CG, Desai RR, Ji Z, Adusumilli S, Jairam N, Ghabra N, Joshi N, Fitch E, Helbig KL, McKnight D, Lindy AS, Zou F, Helbig I, Cooper EC, George AL Jr. High-throughput evaluation of epilepsy-associated KCNQ2 variants reveals functional and pharmacological heterogeneity. JCI Insight. 2022 Mar 8;7(5):e156314. doi: 10.1172/jci.insight.156314. PMID: 35104249; PMCID: PMC8983144.

Wang H, Kunkel DD, Schwartzkroin PA, Tempel BL. Localization of Kv1.1 and Kv1.2, two K channel proteins, to synaptic terminals, somata, and dendrites in the mouse brain. J Neurosci. 1994 Aug;14(8):4588–99. doi:10.1523/JNEUROSCI.14-08-04588.1994. PMID: 8046438; PMCID: PMC6577177.

Wang HS, Pan Z, Shi W, Brown BS, Wymore RS, Cohen IS, Dixon JE, McKinnon D. KCNQ2 and KCNQ3 potassium channel subunits: molecular correlates of the M-channel. Science. 1998 Dec 4;282(5395):1890–3. doi: 10.1126/science.282.5395.1890. PMID: 9836639.

Wang J, Ou SW, Wang YJ. Distribution and function of voltage-gated sodium channels in the nervous system. Channels (Austin). 2017 Nov 2;11(6):534–554. doi: 10.1080/19336950.2017.1380758. Epub 2017 Nov 8. PMID: 28922053; PMCID: PMC5786190

Wang T, Hoekzema K, Vecchio D, Wu H, Sulovari A, Coe BP, Gillentine MA, Wilfert AB, Perez-Jurado LA, Kvarnung M, Sleyp Y, Earl RK, Rosenfeld JA, Geisheker MR, Han L, Du B, Barnett C, Thompson E, Shaw M, Carroll R, Friend K, Catford R, Palmer EE, Zou X, Ou J, Li H, Guo H, Gerdts J, Avola E, Calabrese G, Elia M, Greco D, Lindstrand A, Nordgren A, Anderlid BM, Vandeweyer G, Van Dijck A, Van der Aa N, McKenna B, Hancarova M, Bendova S, Havlovicova M, Malerba G, Bernardina BD, Muglia P, van Haeringen A, Hoffer MJV, Franke B, Cappuccio G, Delatycki M, Lockhart PJ, Manning MA, Liu P, Scheffer IE, Brunetti-Pierri N, Rommelse N, Amaral DG, Santen GWE, Trabetti E, Sedláček Z, Michaelson JJ, Pierce K, Courchesne E, Kooy RF; SPARK Consortium; Nordenskjöld M, Romano C, Peeters H, Bernier RA, Gecz J, Xia K, Eichler EE. Large-scale targeted sequencing identifies risk genes for neurodevelopmental disorders. Nat Commun. 2020 Oct 1;11(1):4932. doi: 10.1038/s41467-020-18723-y. Erratum in: Nat Commun. 2020 Oct 21;11(1):5398. doi: 10.1038/s41467-020-19289-5. PMID: 33004838; PMCID: PMC7530681.

Watanabe K, Kawana E. A horseradish peroxidase study on the mammillothalamic tract in the rat. Acta Anat (basel) 1980;108:394–401. doi: 10.1159/000145322.

Witter MP. Connections of the subiculum of the rat: topography in relation to columnar and laminar organization. Behav Brain Res. 2006;174:251–264. doi: 10.1016/j.bbr.2006.06.022.

Yamagata T, Ogiwara I, Mazaki E, Yanagawa Y, Yamakawa K. Na_V_1.2 is expressed in caudal ganglionic eminence-derived disinhibitory interneurons: Mutually exclusive distributions of Na_V_1.1 and Na_V_1.2. Biochem Biophys Res Commun. 2017 Sep 30;491(4):1070–1076. doi: 10.1016/j.bbrc.2017.08.013. Epub 2017 Aug 4. PMID: 28784306.

Ye J, Tang S, Miao P, Gong Z, Shu Q, Feng J, Li Y. Clinical analysis and functional characterization of KCNQ2-related developmental and epileptic encephalopathy. Front Mol Neurosci. 2023 Jul 11;16:1205265. doi: 10.3389/fnmol.2023.1205265. PMID: 37497102; PMCID: PMC10366601.

Yu H, Rubinstein M, Low MJ. Developmental single-cell transcriptomics of hypothalamic POMC neurons reveal the genetic trajectories of multiple neuropeptidergic phenotypes. Elife. 2022 Jan 19;11:e72883. doi: 10.7554/eLife.72883. PMID: 35044906; PMCID: PMC8806186.

Żakowski W, Braszka Ł, Zawistowski P, Orzeł-Gryglewska J, Jurkowlaniec E. Inactivation of the medial mammillary nucleus attenuates theta rhythm activity in the hippocampus in urethane-anesthetized rats. Neurosci Lett. 2017 Apr 3;645:19–24. doi: 10.1016/j.neulet.2017.02.057. Epub 2017 Feb 22. PMID:28237801.

Żakowski W, Zawistowski P, Braszka Ł, Jurkowlaniec E. The effect of pharmacological inactivation of the mammillary body and anterior thalamic nuclei on hippocampal theta rhythm in urethane-anesthetized rats. Neuroscience. 2017 Oct 24;362:196–205. doi: 10.1016/j.neuroscience.2017.08.043. Epub 2017 Aug 24.PMID: 28844761.

Żakowski W, Zawistowski P. Neurochemistry of the mammillary body. Brain Struct Funct. 2023 Jul;228(6):1379–1398. doi: 10.1007/s00429-023-02673-4. Epub 2023 Jun 28. PMID: 37378855; PMCID: PMC10335970.

Zhang Y, Xue Y, Ma Y, Du X, Lu B, Wang Y, Yan Z. Improved classification and pathogenicity assessment by comprehensive functional studies in a large data set of KCNQ2 variants. Life Sci. 2024 Feb 15;339:122378. doi: 10.1016/j.lfs.2023.122378. Epub 2023 Dec 23. PMID: 38142737.

